# Hastened fusion-dependent endosomal escape improves activity of delivered enzyme cargo

**DOI:** 10.1101/2024.09.27.615476

**Authors:** Angel Luis Vázquez Maldonado, Teresia Chen, Diego Rodriguez, Madeline Zoltek, Alanna Schepartz

## Abstract

There is enormous interest in strategies to efficiently traffic biologics–proteins, nucleic acids, and complexes thereof–into the mammalian cell cytosol and internal organelles. Not only must these materials reach the appropriate cellular locale fully intact and in therapeutically relevant concentrations, they must also retain activity upon arrival. The question of residual activity is especially critical when delivery involves exposure to the late endocytic pathway, whose acidic lumenal environment can denature and/or degrade internalized material. ZF5.3 is a compact, stable, rationally designed mini-protein that efficiently escapes intact from late endocytic vesicles, with or without covalently linked protein cargo. Here, using insights from mechanistic studies on the pathway of endosomal escape and classic principles of zinc(II) bioinorganic chemistry, we re-designed the sequence of ZF5.3 to successfully alter the timing (but not the efficiency) of endosomal escape. The new mini-protein we describe, AV5.3, escapes earlier than ZF5.3 along the endocytic pathway with no loss in efficiency, with or without enzyme cargo. More importantly, earlier endosomal escape translates into higher enzymatic activity of a pH-sensitive enzyme upon arrival in the cytosol. Delivery of the pH-sensitive enzyme dihydrofolate reductase (DHFR) with AV5.3 results in substantial catalytic activity in the cytosol, whereas delivery with ZF5.3 does not. The activity of AV5.3-DHFR upon delivery is sufficient to rescue a genetic DHFR deletion in CHO cells. This work provides evidence that programmed trafficking through the endosomal pathway is a viable strategy for the efficient cytosolic delivery of therapeutic proteins.

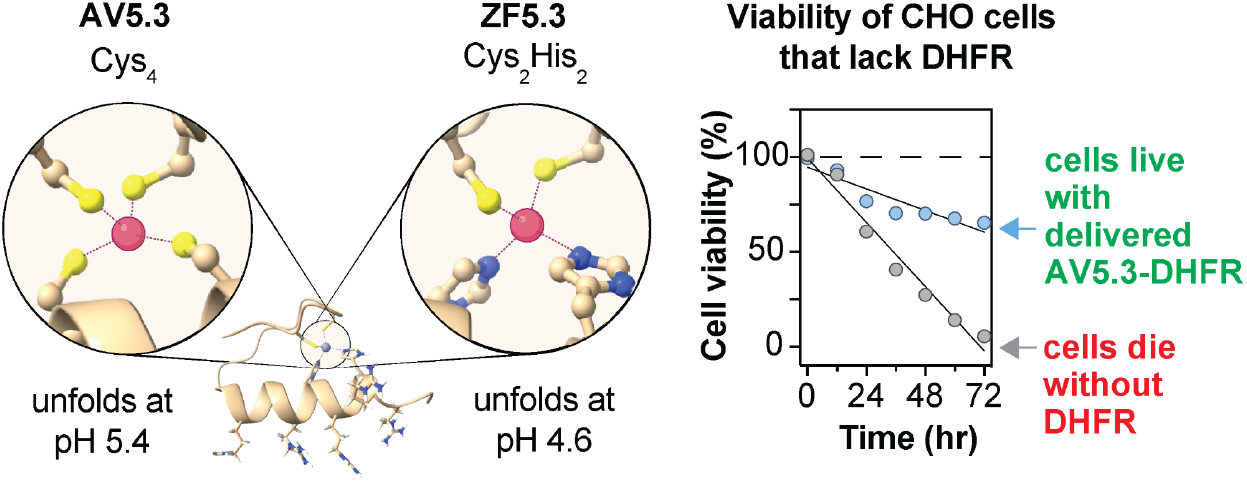

## Introduction

Proteins able to successfully circumnavigate into the mammalian cell cytosol or internal organelles have enormous unrealized potential as replacement enzymes, gene editing tools, protein interaction inhibitors, and bispecific ligands. As a result, multiple potential delivery strategies have been evaluated over the past three decades, including lipid nanoparticles,^1–3^ virus-like particles,^4–7^ super-charged proteins,^8^ charge-masked proteins,^9^ repurposed toxins and nanomachines,^10–12^ as well as literally thousands of molecules described as “cell-penetrating” peptides.^13–16^ Even beyond cell-type specificity, challenges related to both concentration and activity limit all of these approaches. Not only must the delivered cargo reach the appropriate cellular locale fully intact and in therapeutically relevant concentrations, it must also retain activity upon arrival. The question of residual activity is especially critical when delivery involves exposure to the late endocytic pathway, whose acidic lumenal environment denatures and/or degrades internalized materials. It has been estimated that the RNA within FDA-approved SARS-CoV2 vaccines or siRNA therapeutics reach the cytosol with efficiencies significantly less than 10%.^1^ No FDA-approved protein therapeutic acts within the cytosol.

ZF5.3 is a compact, rationally designed mini-protein that delivers proteins to the cytosol or nucleus with high efficiency.^17,18^ Despite its small size (27 amino acids), ZF5.3 is exceptionally stable, with a thermal melting temperature at or above 90°C at pH 7.4. Indeed, ZF5.3 can be isolated intact from the cytosol of treated cells,^17^ and guides multiple classes of proteins, including enzymes,^18–20^ transcription factors,^21^ nanobodies,^22^ and monobodies^20^ into the cytosol and/or nuclei of living cells. In the best cases, the efficiency of delivery to the cytosol reaches or exceeds 50% to establish nuclear or cytosolic concentrations of 500 nM or higher.^17,23,24^ Studies have shown that ZF5.3 escapes into the cytosol from late endolysosomes in a mechanistically distinct process that demands a fully assembled HOPS complex, a ubiquitous tethering machine that promotes late endosomal fusion.^23^ Notably, endosomal escape of ZF5.3 and covalent ZF5.3-conjugates proceeds without leakage of other intralumenal components,^23^ with little or no detectable endosomal damage,^23^ and is especially efficient when the cargo protein is small, intrinsically disordered, or unfolds at a temperature of 35°C or lower.^20^

Recent studies suggest that the unfolding of ZF5.3 itself is also critical for efficient endosomal escape.^24^ Although ZF5.3 is exceptionally stable at pH 7.4, between pH 4 and pH 5 ZF5.3 unfolds cooperatively in a transition initiated by protonation of a single Zn(II)-bound His residue. The p*K*_a_ of this His residue corresponds almost exactly to that of the late endolysosomal lumen, pH 4.6. Evidence that pH-induced unfolding is essential for endosomal escape of ZF5.3 derives from the observation that a ZF5.3 analog that lacks bound Zn(II) and remains folded at low pH is taken up into the endocytic pathway but fails to efficiently reach the cytosol.^24^

Despite the promise of ZF5.3 for cytosolic delivery, the environment within the late endocytic lumen is harsh. The acidic pH, which can be as low as pH 4.5, can degrade RNAs and denature proteins, often irreversibly. Denatured proteins are substrates for lumenal hydrolytic enzymes whose role is to regenerate cellular amino acid building blocks. Although certain therapeutic cargoes successfully delivered by ZF5.3 retain measurable activity after exposure to the endolysosomal lumen, including MeCP2^21^ and a Bcl-11A-targeted bio-protac,^22^ others are likely less robust. We wondered whether we could avoid the detrimental effects of the late endosomal pH by fine-tuning the structure of ZF5.3 to promote escape from an earlier point along the endocytic pathway, that is, a compartment whose pH is higher than 4.6.

Our re-design of ZF5.3 was inspired by the classic bioinorganic chemistry of zinc finger proteins. Like ZF5.3, many zinc finger proteins contain a canonical Cys_2_His_2_ Zn(II) coordination site.^25,26^ But many others, both natural and designed, contain Cys in place of one (Cys_3_His) or both (Cys_4_) His residues.^27^ This change is significant. The p*K*_a_ of a Zn(II)-bound Cys side chain generally falls between 5.0 and 6.0, approximately 2.0 pH units higher than the p*K*_a_ of a Zn(II)-bound His residue.^28^ We therefore hypothesized that variants of ZF5.3 with one or more His-to-Cys substitutions would unfold at pH values higher than 4.6. We hypothesized further that if unfolding is truly critical for endosomal escape, the new variant(s) would escape from endosomal compartments formed earlier along the pathway whose lumen are less acidic. Earlier endosomal escape should translate into improved activity of a pH-sensitive cargo protein.

Here we test these ideas through the design of AV5.3, a variant of ZF5.3 in which Zn(II) is bound not by a Cys_2_His_2_ motif, but instead by a Cys_4_ motif. Like ZF5.3, AV5.3 unfolds cooperatively at low pH, but in this case the pH midpoint occurs at pH 5.4, not 4.6. Despite this difference, AV5.3 and AV5.3-protein complexes are taken up by live cells and traffic into the cytosol with virtually the same efficiency as ZF5.3 and analogous ZF5.3-protein complexes.

With AV5.3, however, cytosolic trafficking depends not only on the activity of the HOPS complex, but also on the activity of the CORVET complex, whose substrates are endosomal vesicles marked by Rab5^29^ and whose lumenal pH is higher - between 6.0 and 6.5.^30^ Finally, we observe that earlier escape is associated with substantially improved activity of a delivered pH-sensitive cargo enzyme. Delivery of the pH-sensitive enzyme dihydrofolate reductase (DHFR) as a conjugate to AV5.3 results in substantially higher cytosolic activity than delivery using ZF5.3. Moreover, only the AV5.3-DHFR conjugate effectively rescues a genetic DHFR knockout in live CHO cells.

## Results

### Chemically tuning the pH required for unfolding: AV5.3 unfolds at a higher pH than ZF5.3

To test whether variants of ZF5.3 containing one or more His-to-Cys substitutions would unfold at pH values higher than 4.6, we prepared two ZF5.3 variants in which the Cys_2_His_2_ Zn(II) coordination site was replaced by either Cys_3_His (CCHC) or Cys_4_ (AV5.3) (Figure 1A-B). CCHC and AV5.3 were synthesized using solid-phase methods, purified by HPLC, and their identities verified using LC-MS (Figure S1). We then used circular dichroism (CD) spectroscopy to assess and compare the pH-dependent secondary structures of CCHC and AV5.3 with that of ZF5.3 (Figure 1C-G). At pH 7.5 and a concentration of 115 µM, the CD spectra of ZF5.3, CCHC, and AV5.3 were qualitatively similar, with pronounced negative ellipticity at 208 nm, as expected for a molecule containing a ββα zinc finger fold^31,32^ (Figure 1C). As found for ZF5.3,^24^ these features depend on the presence of Zn(II) (Figure S2) but are independent of temperature between 5 and 95°C (Figure 1D).

**Figure 1.**
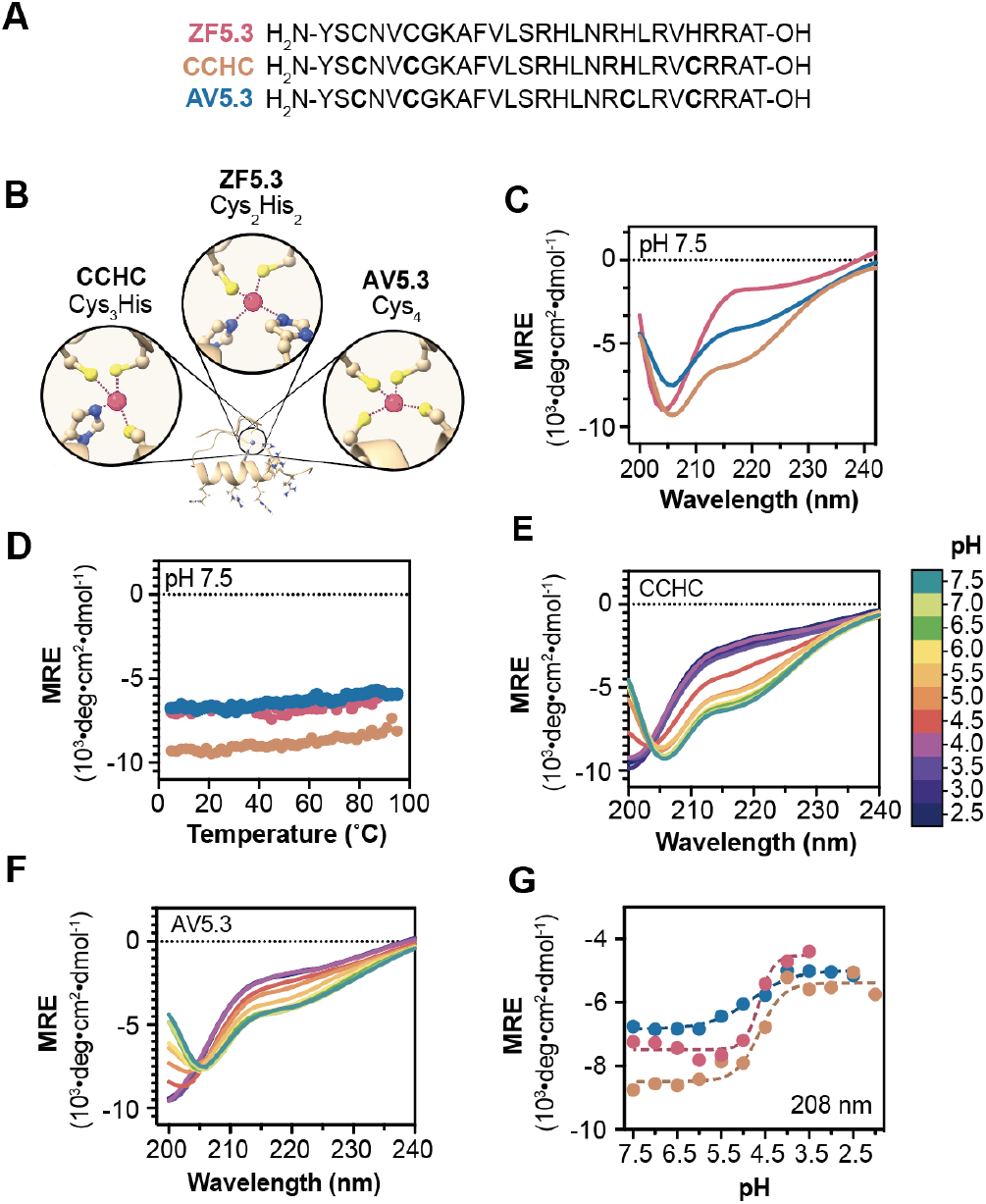
pH-induced unfolding of ZF5.3 variants AV5.3 and CCHC. **(A)** Primary sequences of ZF5.3, CCHC, and AV5.3, in which one (His_23_) or both (His_19_ and His_23_) Zn(II)-coordinating His ligands are substituted by Cys. **(B)** Illustration that highlights the differences between ZF5.3, CCHC, and AV5.3’s Zn(II) coordination-site. **(C)** Smoothed wavelength-dependent and **(D)** temperature-dependent CD spectra of ZF5.3, CCHC, and AV5.3 at a concentration of 115 µM and pH 7.5 in a buffer composed of 20 mM Tris-HCl, 150 mM KCl. Panel **(C)** shows a plot of the mean residue ellipticity at 208 nm as the temperature was increased from 2°C to 95°C every 2°C. Smoothed wavelength-dependent CD spectra of **(E)** CCHC or **(F)** AV5.3, both at a concentration of 115 µM, monitored at every 0.5 pH unit between pH 7.5 and 2.5. For details, see **Supplementary Information. (G)** Plot of the mean residue ellipticity at 208 nm as the pH of a 115 µM solution of ZF5.3, CCHC, or AV5.3 is lowered from pH 7.5 to pH 2.5. These data were fitted using a Boltzmann sigmoid dose response (R^2^ ≥ 0.95) using GraphPad Prism 10 software.

To evaluate the influence of pH on secondary structure, CD spectra were recorded at pH values from 2.5 to 7.5 (Figure 1E-F). An overlay of the spectra for CCHC and AV5.3 revealed shifts in the primary ellipticity minimum towards shorter wavelengths as the pH decreased from 7.5 to 2.5. Similar changes were previously reported for ZF5.3 and are consistent with a pH-induced loss of structure that was confirmed using NMR.^24^ A plot depicting the change in ellipticity at 208 nm as a function of pH revealed cooperative transitions for both ZF5.3 and CCHC with a transition over 2 pH units, whereas the transition for AV5.3 was broader. The transition midpoint occurred at pH 4.6 for CCHC and ZF5.3, whereas the transition midpoint for AV5.3 was centered at 5.4, almost a full pH unit higher (Figure 1F). These findings confirm that the Zn(II) coordination sphere can be tuned to alter the unfolding pH of molecules related to ZF5.3 and that AV5.3 undergoes a pH- and Zn(II)-dependent structural transition with a mid-point that is substantially higher than ZF5.3. CCHC was not studied further because its pH-dependent unfolding transition was virtually identical to that of ZF5.3. Notably, the unfolding pH of AV5.3 (5.4) corresponds most closely to that of the late endosomal lumen (approximately 5.5), whereas that of ZF5.3 (4.6) is closer to the pH of a lysosome.^33^

### AV5.3 traffics efficiently into the cytosol of Saos-2 cells

Next we used confocal microscopy and flow cytometry (FC) to evaluate whether the difference in pH-dependent unfolding of ZF5.3 and AV5.3 was accompanied by a change in overall cell uptake. Variants of ZF5.3 and AV5.3 carrying an *N*-terminal *N*^ε^-azido-L-lysine residue were prepared by solid phase peptide synthesis and fluorescently labeled upon reaction with DBCO-functionalized lissamine rhodamine B to generate AV5.3^Rho^ and ZF5.3^Rho^ (see Figure S1 and **Supplementary Information**). Following purification, AV5.3^Rho^ and ZF5.3^Rho^ were added at concentrations between 0.1 and 1 μM to human osteosarcoma (Saos-2) cells and incubated for 30 min. The cells were washed with trypsin to eliminate surface-bound protein and visualized using confocal microscopy.

Confocal microscopy images of Saos-2 cells treated with either AV5.3^Rho^ or ZF5.3^Rho^ show clear evidence of uptake, with substantial punctate fluorescence distributed throughout the cell interior (Figure 2A). Quantification of the internalized fluorescence as a function of AV5.3^Rho^ or ZF5.3^Rho^ concentration using flow cytometry indicated that ZF5.3^Rho^ is taken up slightly more efficiently than AV5.3^Rho^. On average, cells treated with ZF5.3^Rho^ showed 2-4-fold higher fluorescence at all treatment concentrations relative to cells treated with AV5.3^Rho^, with larger differences at higher treatment concentrations (Figure 2B). In both cases, whether assessed qualitatively using confocal microscopy (Figure 2A) or quantitatively using flow cytometry (Figure 2B), the uptake of fluorescence was dose-dependent. Overall these studies indicate that conversion of the Cys_2_His_2_ coordination sphere in ZF5.3 to the Cys_4_ coordination sphere in AV5.3 has a small but measurable effect on overall uptake by Saos-2 cells.

**Figure 2.**
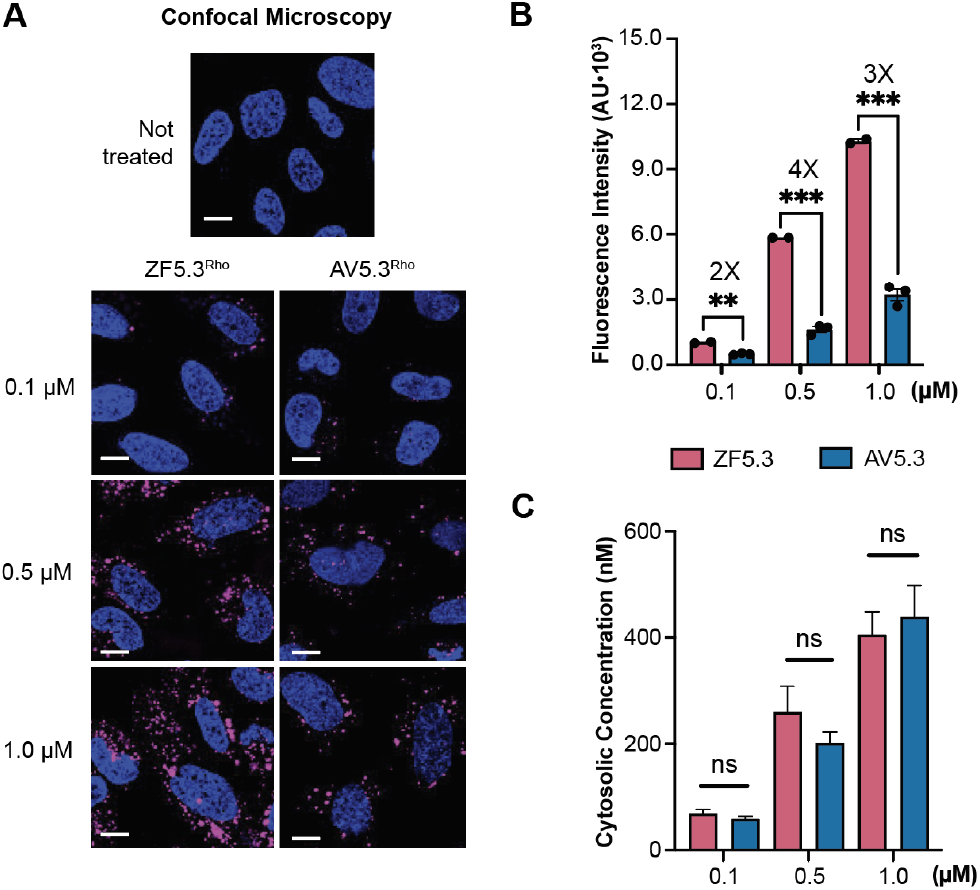
AV5.3^Rho^ reaches the cytosol as well as ZF5.3^Rho^, despite lower uptake. **(A)** 2D confocal microscopy images of Saos-2 cells incubated with the indicated concentration of ZF5.3^Rho^ or AV5.3^Rho^. Scale bar: 10 μm. Plots showing **(B)** flow cytometry (FC) analysis of total cellular uptake; or **(C)** fluorescence correlation spectroscopy (FCS) analysis to establish the cytosolic concentrations of ZF5.3^Rho^ and AV5.3^Rho^ achieved after a 30 min incubation. FC values are provided as median fluorescence intensity (MFI) for readings of lissamine rhodamine B excitation using the yellow laser 1 (585/16 filter); *n* = 20 000 total cells per biological replicate. Each condition contains at least two biological replicates each (median ± SEM). For FCS data, *n* > 20 for each condition with two biological replicates each (median ± SEM). Statistical significance comparing the given concentrations was assessed using the Brown–Forsythe and Welch one-way analysis of variance (ANOVA) followed by an unpaired *t* test with Welch’s correction. \*\*\*\**p* ≤ 0.0001, \*\*\**p* ≤ 0.001, \*\**p* ≤ 0.01, \**p*≤ 0.05.

We next made use of fluorescence correlation spectroscopy (FCS) to assess what fraction of the material taken up into the endocytic pathway trafficked successfully into the cytosol. FCS is a unique tool for assessing endosomal escape, as it provides both the precise concentration of a fluorescent molecule in the cytosol as well as its diffusion constant.^34^ We measured the *in vitro* diffusion constants of ZF5.3^Rho^ and AV5.3^Rho^ to determine the cut-offs for an appropriate range of diffusion constants measured *in cellula* by FCS (Figure S3). FCS analysis of Saos-2 cells treated for 30 min with AV5.3^Rho^ or ZF5.3^Rho^ revealed that both molecules reached the cytosol efficiently and in a dose-dependent manner (Figure 2C). Treatment of Saos-2 cells with 0.1, 0.5, and 1 μM AV5.3^Rho^ led to average cytosolic concentrations of 59 (± 5), 201 (± 21), and 439 (± 58) nM, respectively, corresponding to delivery efficiencies between 40% and 59%.

In comparison, treatment of Saos-2 cells with analogous concentrations of ZF5.3^Rho^ led to average cytosolic concentrations of 69 (± 8), 260 (± 49), and 406 (± 43) nM, corresponding to delivery efficiencies between 41% and 69%. Thus we found no statistically significant difference between the cytosolic concentrations established in Saos-2 cells treated with equivalent concentrations of AV5.3^Rho^ or ZF5.3^Rho^, despite the higher uptake of ZF5.3 detected using flow cytometry (Figure 2B). Our results demonstrate that, despite slightly lower overall uptake, AV5.3 reached the cytosol as well as ZF5.3. This result implies that AV5.3 escapes from the endocytic pathway with an efficiency that is at least as high as ZF5.3.

One of the unique attributes of ZF5.3 is its ability to escape from endosomes without detectable endosomal rupture; co-treating cells with ZF5.3 and Lys9^Rho^ under conditions that result in high cytosolic concentrations of ZF5.3 led to no detectable cytosolic fluorescence due to Lys9^Rho^.^23^ To evaluate whether this behavior was also recapitulated by AV5.3, we co-treated cells with AV5.3 and Lys9^Rho^ and evaluated the cells using FCS (Figure S4). We detected no no evidence for Lys9^Rho^ in the cytosol when added together with AV5.3.

To further characterize the trafficking of AV5.3, also compared its localization within Rab5-, Rab7- and Lamp1-positive vesicles using validated GFP markers for these three organelles. These studies revealed high co-localization of AV5.3^Rho^ with Rab7- and Lamp1-positive vesicles, with minimal localization within Rab5-positive vesicles (Figure S5). This pattern is almost identical to that of ZF5.3^Rho^.^23^

### AV5.3 reaches the cytosol in a CORVET- and HOPS-dependent manner

Previous research has shown that both ZF5.3 and ZF5.3-protein conjugates rely on the HOPS complex for cytosolic access.^20,21,23,35–37^ When cells are depleted of the Rab7-binding HOPS subunits VPS39 and VPS41, the ability of ZF5.3 to reach the cytosol is substantially diminished. The same is true for ZF5.3-protein conjugates that efficiently reach the cytosol.^20,21,23^ In contrast, knockdown of the Rab5-binding CORVET subunits VPS8 and TGF-BRAP1 fails to diminish the ability of ZF5.3 to reach the cytosol. In fact, in some cases, depletion of TGF-BRAP1 improves cytosolic delivery of ZF5.3 and ZF5.3-protein conjugates,^20,21,23^ perhaps because it increases the intracellular concentration of HOPS.^38^ We used analogous siRNA experiments to evaluate the effect of HOPS- and CORVET depletion on the delivery efficiency of AV5.3. If the higher-pH unfolding of AV5.3 leads to escape from an earlier, high-pH endocytic compartment, then escape of AV5.3 should show an higher dependence on CORVET, which operates at an earlier stage of the endocytic pathway.^33,39^

Saos-2 cells were transfected with siRNAs targeting each of the CORVET-specific subunits VPS8 and TGF-BRAP1 or the HOPS-specific subunits VPS39 and VPS41; knockdown efficiencies were established using quantitative PCR (Figure S6). After siRNA transfection, Saos-2 cells were treated with 1 µM AV5.3^Rho^ or ZF5.3^Rho^ for 30 min and the cells analyzed using confocal microscopy, flow cytometry, and FCS (Figure 3).

**Figure 3.**
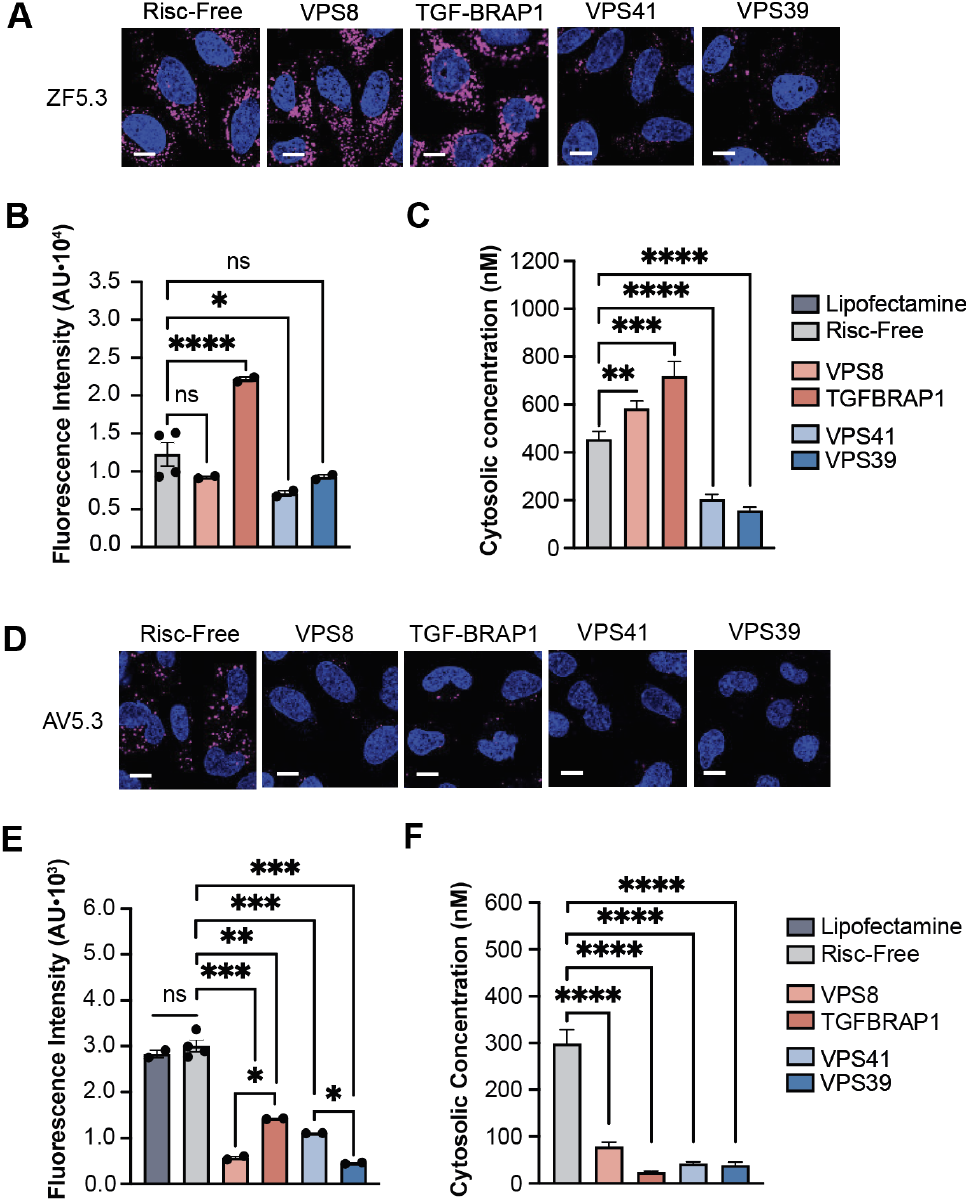
AV5.3 reaches the cytosol in a CORVET- and HOPS-dependent manner. **(A)** 2D confocal microscopy images of Saos-2 cells transfected with siRNAs targeting CORVET (VPS8 and TGF-BRAP1) or HOPS (VPS39 and VPS41) and incubated with ZF5.3^Rho^. For details, see **Supplementary Information.** Scale bar: 10 μm. **(B-C)** Plots illustrating the effects of siRNAs targeting CORVET (VPS8 and TGF-BRAP1) and HOPS (VPS39 and VPS41) on the total uptake (flow cytometry, median fluorescence intensity) and cytosolic access (FCS, nM) of 1.0 µM ZF5.3. **(D)** 2D confocal microscopy images of Saos-2 cells transfected with siRNAs targeting CORVET (VPS8 and TGF-BRAP1) or HOPS (VPS39 and VPS41) and incubated with AV5.3^Rho^. For details, see **Supplementary Information**. Scale bar: 10 μm. **(E-F)** Plots illustrating the effects of siRNAs targeting CORVET (VPS8 and TGF-BRAP1) and HOPS (VPS39 and VPS41) on the total uptake (flow cytometry, median fluorescence intensity) and cytosolic access (FCS, nM) of 1.0 µM AV5.3. The data shown represents at least two biological replicates; *n* = 20 000 per condition in total for flow cytometry and *n* > 15 per condition for FCS. Error bars represent the SEM (\*\*\*\**P* < 0.0001, \*\*\**P* < 0.001, \*\**P* < 0.01, \**P* < 0.05, and not significant (ns) for *P* > 0.05) from one-way ANOVA with unpaired *t* test with Welch’s correction.

Cells treated with ZF5.3^Rho^ responded to HOPS and CORVET depletion in a manner consistent with previous reports.^23^ Depletion of HOPS-specific subunits VPS39 and VPS41 visually decreased the punctate fluorescence evident by confocal microscopy (Figure 3A), reduced by 33% the level of internalized ZF5.3^Rho^ fluorescence detected by flow cytometry (Figure 3B), and reduced by 60% the concentration of ZF5.3^Rho^ that reached the cytosol as determined by FCS (Figure 3C). Depletion of CORVET-specific subunits also had the expected effects (Figure 3B,C).

Cells treated with AV5.3^Rho^ responded differently to HOPS and CORVET depletion than cells treated with ZF5.3. In this case, depletion of either HOPS subunits (VPS39, VPS41) or CORVET subunits (VPS8, TGF-BRAP1) decrease the overall uptake of AV5.3^Rho^ (Figure 3D) and its localization to the cytosol (Figure 3E). Depletion of HOPS subunits VPS39 and VPS41 decreased the concentration of AV5.3^Rho^ that reached the cytosol by 86 and 87%, respectively, while depletion of CORVET subunits VPS8 and TGF-BRAP1 led to a decrease of 74 and 92%, respectively. No changes in the cytosolic localization of AV5.3^Rho^ were observed when cells were mock-transfected or transfected with a chemically modified siRNA that fails to engage with the Risc complex (Risc-free). It is notable that the cytosolic localization of AV5.3 is almost completely abolished by either HOPS or CORVET knockdown, suggesting an interplay between these two tethering complexes that is not fully understood. Regardless, these results indicate that both CORVET and HOPS contribute to the cytosolic localization of AV5.3, and that AV5.3 escapes, at least in part, from Rab5+ vesicles that are substrates for CORVET. Moreover, the dependence of AV5.3 delivery on both HOPS and CORVET provides additional support for a mechanistic link between ZF5.3/AV5.3 unfolding and endosomal escape.^24^

### AV5.3 provides DHFR with an alternate but equally effective path into the cytosol

Next we sought to determine whether the alternative, HOPS and CORVET-dependent path into the cytosol taken by AV5.3 also supports the delivery of protein cargo. One of the most efficiently delivered cargos when fused to ZF5.3 is dihydrofolate reductase (DHFR), in large part because the DHFR T_M_ is low in the absence of bound ligand.^40^ Samples of AV5.3–DHFR and ZF5.3–DHFR were purified to homogeneity from *E. coli* and characterized using LC/MS and CD (Figure S7 and S8). The effects of pH and the presence of DHFR’s ligand, methotrexate (MTX), on the secondary structure and thermal stability of AV5.3–DHFR and ZF5.3–DHFR were virtually identical (Figure S8). Rhodamine-labeled derivatives of each conjugate (AV5.3–DHFR^Rho^ and ZF5.3–DHFR^Rho^) were prepared using sortase as previously described (Figure S7).^20^

To evaluate the delivery of protein cargo, Saos-2 cells were treated with between 0.1 and 1 μM AV5.3-DHFR^Rho^ or ZF5.3–DHFR^Rho^ for 1 h and visualized using confocal microscopy, FC, and FCS as described above. Confocal microscopy and FC revealed that AV5.3-DHFR^Rho^ and ZF5.3–DHFR^Rho^ were taken up almost identically by Saos-2 cells and in a dose-dependent manner (Figure 4A, B). The overall uptake of ZF5.3–DHFR^Rho^ by Saos-2 cells is comparable to levels observed previously.^20^ In a similar way, FCS analysis revealed that both AV5.3-DHFR^Rho^ or ZF5.3–DHFR^Rho^ reached the Saos-2 cytosol in a dose-dependent manner and with almost identical efficiency (Figure 4C). These delivery efficiencies were confirmed upon Western blot analysis of isolated cytosolic fractions (Figure S9). Finally, knockdown experiments revealed the overall uptake of AV5.3-DHFR as well as its ability to reach the cytosol depends on both the Rab7-binding subunits of HOPS as well as the Rab5-binding components of CORVET (Figure 4E). These results confirm that the alternative, HOPS and CORVET-dependent path into the cytosol taken by AV5.3 fully supports the delivery of protein cargo.

**Figure 4.**
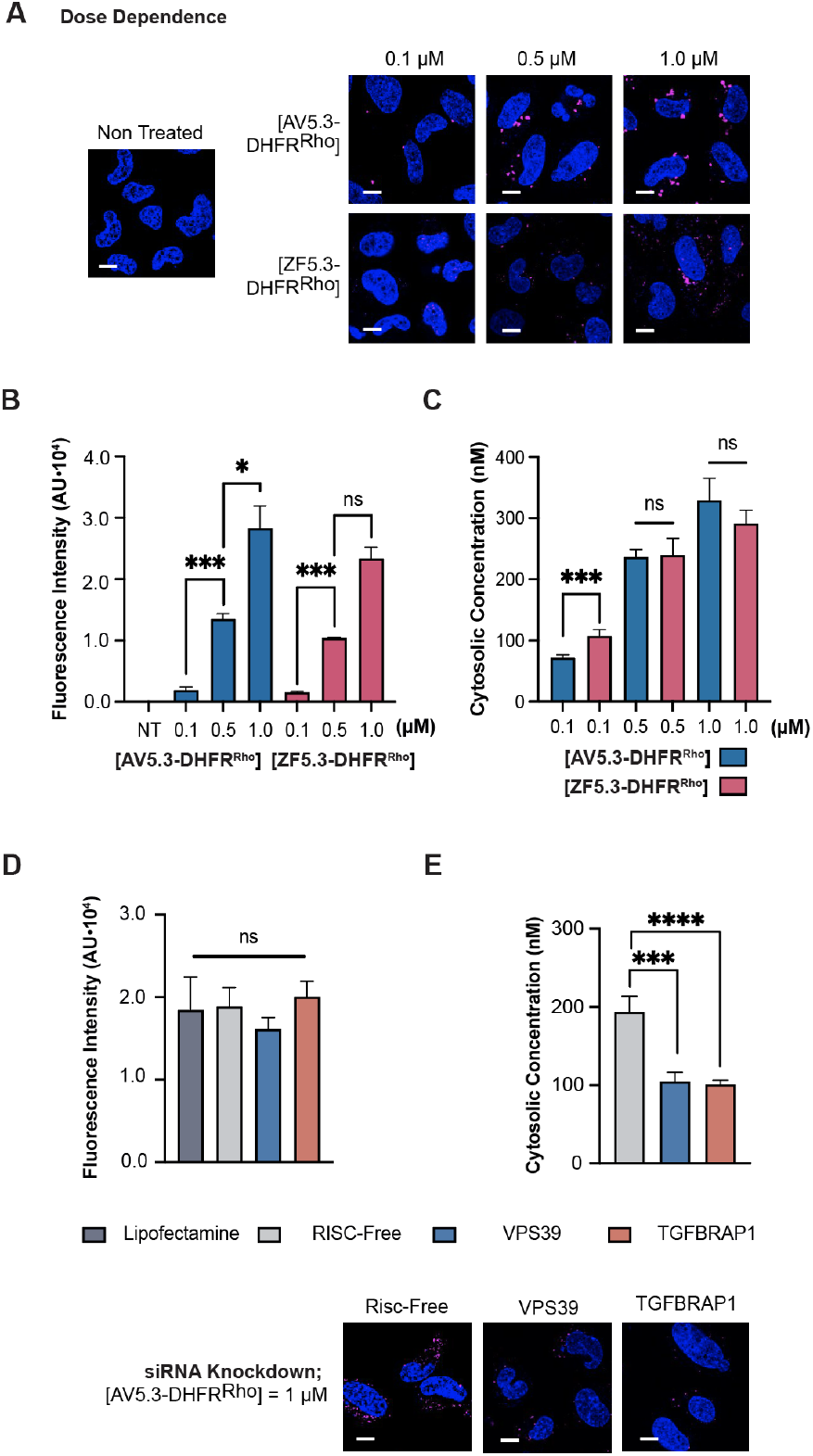
DHFR reaches the cytosol in a CORVET- and HOPS-dependent manner when delivered by AV5.3. **(A)** 2D confocal microscopy images of Saos-2 cells incubated with the indicated concentration AV5.3-DHFR^Rho^ as described in **Supplementary Information.** Scale bar: 10 μm. **(B-C)** Plots illustrating the dose dependence of AV5.3-DHFR with respect to both total uptake (flow cytometry, median fluorescence intensity) and delivery to cytosol (FCS, nM). **(D-E)** 2D confocal microscopy images and plots illustrating the effects of CORVET (TGF-BRAP1) and HOPS (VPS39) knockdowns on the total uptake of AV5.3-DHFR (flow cytometry, median fluorescence intensity) and its ability to reach the cytosol (FCS, nM). At least two biological replicates were performed for each experiment; *n* = 20 000 per condition in total for flow cytometry and *n* > 15 per condition for FCS. Error bars represent the SEM (\*\*\*\**P* < 0.0001, \*\*\**P* < 0.001, \*\**P* < 0.01, \**P* < 0.05, and not significant (ns) for *P* > 0.05) from one-way ANOVA with unpaired *t* test with Welch’s correction.

### Comparing the enzymatic activity of delivered AV5.3-DHFR and ZF5.3-DHFR: in vitro controls

The data presented above support a model in which AV5.3 escapes from the endocytic pathway into the cytosol, at least in part, from earlier, and presumably less acidic endocytic compartments than does ZF5.3. This difference should improve the residual activity of delivered proteins or enzymes that struggle to refold and/or regain activity after exposure to low pH. Mammalian DHFR is one such enzyme. Although DHFR refolds after guanidinium hydrochloride-induced denaturation,^41^ it fails to refold after heat treatment or exposure to low pH.^20^

To evaluate the residual activities of ZF5.3–DHFR and AV5.3–DHFR post-delivery, we first assessed their catalytic activities *in vitro* in comparison with a human DHFR standard. DHFR catalyzes the reduction of 7,8-dihydrofolate (DHF) to 5,6,7,8-tetrahydrofolate (THF) using a single equivalent of NADPH as a cofactor (Figure 5A). Its activity is conveniently measured spectrophotometrically by monitoring the decrease in NADPH absorbance at 340 nm in a reaction mixture supplemented with enzyme and DHF (Figure 5B). To evaluate enzyme activities in vitro, solutions of hDHFR, ZF5.3-DHFR, or AV5.3-DHFR at 250 nM were prepared and the enzymatic reaction initiated upon addition of 50 µM DHF.^42^ The resulting decrease in A_340_ was monitored as a function of time (see **Supplementary Information** and Figure S11) and used to calculate specific activity in units of µmole/min/mg protein.

**Figure 5.**
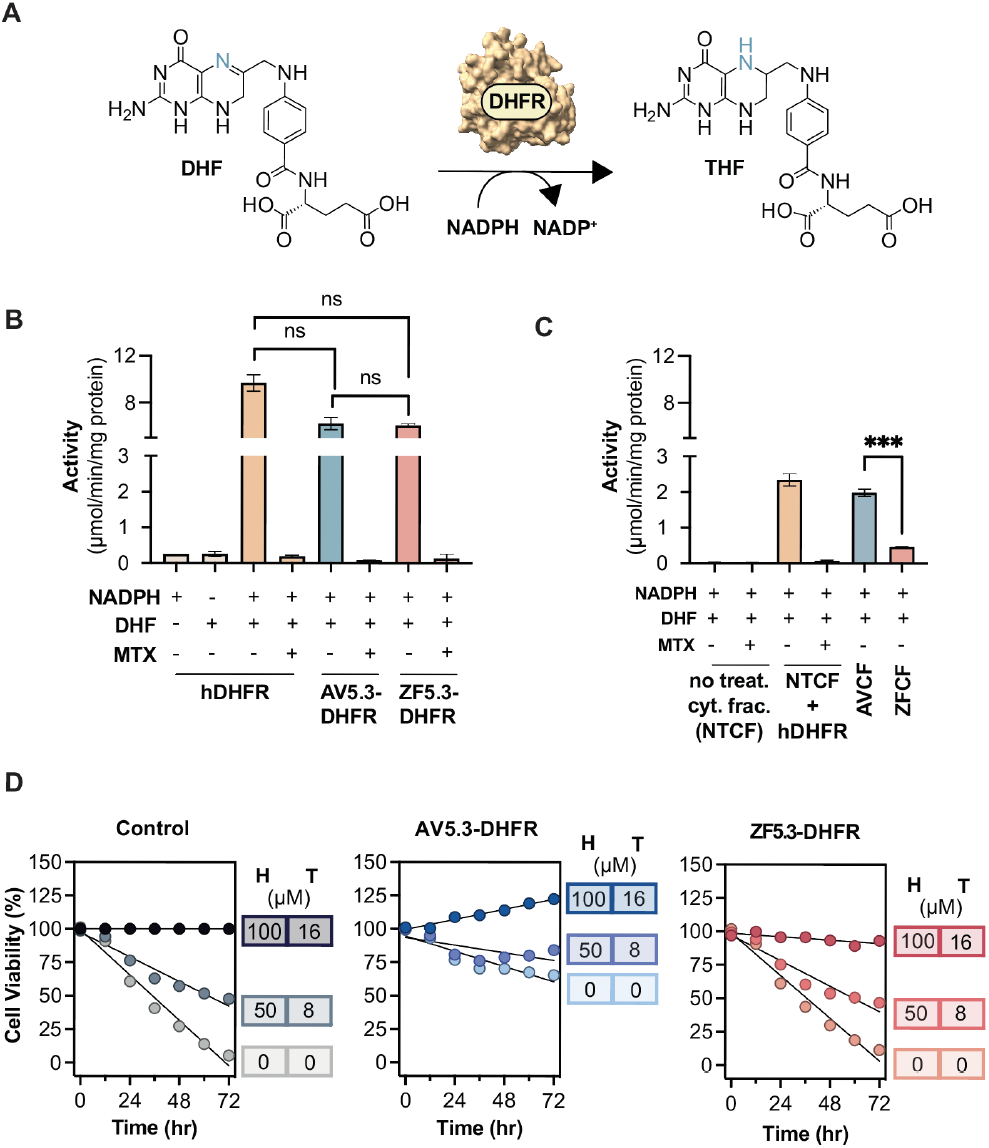
DHFR enzyme retains activity when delivered by AV5.3. **(A)** DHFR catalyzes the reduction of 7,8-dihydrofolate (DHF) to 5,6,7,8-tetra-hydrofolate (THF) utilizing NADPH as cofactor. **(B)** Plot comparing the specific activity of hDHFR, AV5.3-DHFR and ZF5.3-DHFR. All three enzymes are inhibited completely by methotrexate (MTX). **(C)** Specific activity values of isolated cytosol extracts from non-treated CHO/dhFR-cells (NTCF) in the absence or presence of 200 nM hDHFR, alongside those from CHO/dhFR-cells treated with 1 µM AV5.3-DHFR (termed AVCF) or 1 µM ZF5.3-DHFR (termed ZFCF). **(D)** DHFR rescue assay monitoring the time-dependent viability of CHO/dhFr-cells in the presence of varying concentrations of hypoxanthine (H) and thymidine (T) supplement and in the absence (control) or presence of 1 µM AV5.3-DHFR or 1 µM ZF5.3-DHFR. Cell viability was monitored every 12 h for 72 h using CellTiter-Glo® 2.0 Cell Viability Assay and normalized to the viability of CHO/dhFr-cells supplemented with 100 µM hypoxanthine and 16 µM thymidine.

Analysis of the time-dependent decreases in NADPH absorbance revealed that both ZF5.3-DHFR and AV5.3-DHFR are catalytically active (Figure 5C). The activity of all three proteins fell between 6.1 and 9.7 moles/min/mg protein. As expected, pre-incubation of hDHFR, AV5.3-DHFR or ZF5.3-DHFR with 2 molar equivalents of methotrexate (MTX) led to a complete loss of enzymatic activity (Figure 5C). Thus, when measured in vitro, there was no significant difference between the specific activities of AV5.3-DHFR and ZF5.3-DHFR and no significant difference between either of these two conjugates and hDHFR itself.

### Cytosolic DHFR is more catalytically active when delivered by AV5.3

To evaluate the activities of cytosolic ZF5.3-DHFR and AV5.3-DHFR post-delivery, we made use of a commercial CHO cell line that lacks DHFR (CHO/dhFr-). Although DHFR is otherwise essential, CHO/dhFr-cells remain viable and grow upon addition of hypoxanthine and thymidine to compensate for the absence of endogenous DHFR. We prepared cytosolic extracts of CHO/dhFr-cells both before and after treatment with ZF5.3-DHFR or AV5.3-DHFR, and assessed residual DHFR activity as described above (Figure 5D). Cytosolic extracts of CHO/dhFr-that had not been treated with ZF5.3-DHFR or AV5.3-DHFR showed no detectable DHFR activity, as expected. Supplementing these non-treated extracts with 250 nM hDHFR restored DHFR activity to a value of 2.34 moles/min/mg protein; this activity was abolished in the presence of 500 nM MTX (Figure 5D).

Next we treated live cultures of CHO/dhFr-cells with 1 µM of either AV5.3-DHFR or ZF5.3-DHFR for 1 h, prepared cytosolic extracts, and tested the extracts for DHFR activity. Cytosolic extracts of CHO/dhFr-cells treated with 1 µM AV5.3-DHFR were characterized by a DHFR activity of 2.0 moles/min/mg, 15% lower than the activity observed when untreated extracts were supplemented with 250 nM hDHFR. This value is only slightly lower than the concentration of AV5.3-DHFR that reaches the Saos-2 cytosol after a 1 µM treatment (329.1 ± 36 nM, see Figure 4). This result suggests that AV5.3-DHFR retains significant activity even after exposure to the endocytic pathway. In contrast, cytosolic extracts of CHO/dhFr-cells treated with 1 µM ZF5.3-DHFR were characterized by a DHFR activity of 0.5 moles/min/mg, a value that is roughly 80% lower than the activity observed when untreated extracts were supplemented with 250 nM hDHFR, and equally lower than the concentration of ZF5.3-DHFR that reaches the Saos-2 cytosol after a 1 µM treatment (291.2 ± 22 nM). The substantial difference between the residual activities of AV5.3-DHFR or ZF5.3-DHFR in cytosolic extracts post-delivery provides direct evidence that AV5.3 not only delivers protein cargos efficiently but also provides confidence that even highly pH-sensitive cargo proteins and enzymes will retain activity upon reaching the cytosol.

### AV5.3-DHFR rescues the DHFR deficiency of CHO/dhFR-cells

Finally, we asked whether AV5.3-DHFR would rescue, in full or in part, the DHFR deficiency of CHO/dhFR-cells. CHO/dhFR-cells fail to grow without addition of hypoxanthine and thymidine, and their viability, as measured by ATP activity, diminishes slowly over the course of 72 h. Supplementation every 12 h with 100 µM hypoxanthine and 16 µM thymidine fully maintains cell viability as measured by the concentration of ATP (CellTiter-Glo® 2.0 Cell Viability Assay). When the concentration of hypoxanthine and thymidine was reduced by half, cell viability decreases by roughly 50% over 48 h (Figure 5D). Treatment of CHO/dhFR-cells with 1 µM ZF5.3-DHFR had no significant effect on cell viability in the presence or absence of hypoxanthine and thymidine supplement. In contrast, CHO/dhFR-cells treated with 1 µM AV5.3-DHFR remained viable over the course of 72 h in the presence or absence of hypoxanthine and thymidine supplement. In the absence of any supplement, viability of AV5.3-DHFR-treated cells after 72 h was roughly 35% lower than the viability of untreated CHO/dhFR-in the presence of complete hypoxanthine and thymidine supplement. In the presence of half strength supplements, cells treated with 1 µM AV5.3-DHFR were only 16% less viable than untreated CHO/dhFR-in the presence of complete hypoxanthine and thymidine supplements. In the presence of full strength hypoxanthine and thymidine supplement, CHO/dhFR-cells treated with 1 µM AV5.3-DHFR showed almost 25% greater proliferation than cells lacking AV5.3-DHFR treatment. We conclude from these data that the residual enzyme activity of AV5.3-DHFR upon delivery to the cytosol is sufficient to rescue a genetic DHFR deletion in CHO cells.

## Summary

This project was initiated to overcome one of the major challenges facing any macromolecule delivery strategy that relies on the endocytic pathway: exposure of cargo to low pH. Using insights derived from the mechanism and pH-dependence of fusion-dependent endosomal escape, we re-designed the sequence of a mini-protein that reaches the cytosol efficiently to hasten the timing of endosomal escape. When this new mini-protein, AV5.3, is conjugated to an acid-labile enzyme cargo, dihydrofolate reductase (DHFR), delivery efficiency is unaffected but the residual activity of DHFR in the cytosol is substantially improved. This work provides evidence that fusion-dependent endosomal escape can be fine-tuned to improve the residual activity of proteins that rely on endocytic trafficking to reach the cytosol and represents a viable strategy for efficient cytosolic delivery of therapeutic proteins. It also provides *de facto* support that endosomal escape of mini-proteins like ZF5.3 and AV5.3 demand protein unfolding in acidic endosomal compartments.^24^

## ASSOCIATED CONTENT

### Supporting Information

Supporting information includes supplementary methods, figures, and tables. The Supporting Information is available free of charge on the ACS Publications website.

## AUTHOR CONTRIBUTIONS

This project was conceived and developed by A.V.M. and A.S. A.V.M. performed the experiments described in Figures 1-5; T.C. performed qPCR analyses in Figure S6; D.R. assisted with the acquisition of data in Figures S7 and S8; M.Z. provided purified samples of ZF5.3-DHFR. A.V.M. and A.S. analyzed the data and wrote the manuscript with input from all authors.

## ACKNOWLEDGEMENT

Research reported in this publication was supported by the National Institute of General Medical Sciences of the National Institutes of Health under Award R35GM134963. It was also supported by the National Science Foundation (CHE-2203903 to A.S.).We are grateful to all members of the Schepartz lab for support and helpful discussions.

