## Supplementary Material for "Hastened fusion-dependent endosomal escape improves activity of delivered enzyme cargo"

##### Table of Contents

I. Reagents and Chemicals 3
II. Solid-phase peptide synthesis 4
  A. Preparation of ZF5.3, CCHC, and AV5.3 4
  B. Preparation of ZF5.3Rho and AV5.3$^{Rho}$ 5
  C. Reconstitution of ZF5.3 and AV5.3 from stock solutions 5
III. Circular dichroism 5
  A. Instrumentation 5
  B. pH-dependent assays 5
  C. Temperature-dependent assays 6
IV. Plasmids 6
  A. Preparation of pAV-DHFR 6
  B. Preparation of pZF-DHFR 7
V. Protein Expression, Purification, and Characterization 7
  A. AV5.3-DHFR 7
    Expression 7
    Purification 7
    Characterization 8
  B. ZF5.3-DHFR 8
VI. Rhodamine labeling of proteins 8
  A. AV5.3-DHFR$^{Rho}$ 8
    Sortase reaction of AV5.3-DHFR and purification 8
    Characterization of AV5.3-DHFR$^{Rho}$ 8
  B. ZF5.3-DHFRRho 9
VII. Delivery methods 9
  A. Cell culture 9
  B. Delivery procedure 9
  C. Confocal microscopy 10
    Colocalization Experiment 10
  D. Flow cytometry 10
  E. Fluorescence correlation spectroscopy (FCS) 10
  F. FCS data analysis 11
VIII. Endosomal leakage assay 12
IX. siRNA knockdowns 13

A. Knockdown of Saos-2 cells for delivery experiments 13
B. siRNA knockdown validation by qPCR 13
X. DHFR activity assays 13
A. In vitro sample preparation 13
B. Cytosolic fractionation method 13
Cell harvesting 14
Cell lysis and fractionation 14
Western Blot 14
Cytosolic fraction sample preparation for activity assay 14
C. DHFR activity assay 14
D. DHFR rescue assay 15
XI. Statistics 15
XII. Supplementary Tables 1-3 16
Table S1: Relevant Peptide, Protein, and DNA/RNA Sequences 16
Table S2: Summary Values for Intracellular Delivery Experiments 18
Table S3: Statistical information for all comparisons 20
XIII. Supplementary Figures 1-12 25
Figure S1. LC-MS of purified ZF5.3, CCHC, and AV5.3 and Rho-labeled derivatives.. 25
Figure S2. Wavelength-dependent CD of apo and Zn(II)-coordinated AV5.3 and CCHC. 26
Figure S3. Autocorrelation traces generated during FCS *in vitro* and in Saos-2 cells 27
Figure S4. Lys9$^{Rho}$ leakage assay.. 28
Figure S5: AV5.3$^{Rho}$ accumulates in Rab7+ and Lamp1+ endosomes.. 29
Figure S6. qPCR analysis of siRNA knockdowns.. 30
Figure S7. LC-MS analysis showing DHFR proteins and Rho-labeled derivatives.. 31
Figure S8. Wavelength-dependent CD spectra and temperature melts AV5.3-DHFR... 32
Figure S9. AV5.3-DHFR is fully intact when isolated from the cytosol of Saos-2 cells.. 33
Figure S10. Effect of MTX on the total uptake and cytosolic delivery of AV5.3-DHFR$^{Rho}$.. 34
Figure S11. DHFR proteins activity assay curves.. 35
Figure S12. DHFR protein rescue in CHO/dhFr- cells.. 36
XIII. References 37

#### I. Reagents and Chemicals

Chemicals

Isopropyl β-D-thiogalactopyranoside (# 367-93-1) (American Bio), Calcium Chloride (#C5670), Fmoc-Gly-OH (#47627), Fmoc-Lys(Mtt)-OH (#852065), Lissamine Rhodamine B Sulfonyl Chloride (#86186), Lissamine™ Rhodamine B Ethylenediamine (#L2424) (Thermo Fisher Scientific), DBCO-NHS Ester (#CCT-1491) (VWR), N,N-Diisopropylethylamine (#387649), Hexafluorophosphate Benzotriazole Tetramethyl Uronium (#AS-21015) (Anaspec), 1-Hydroxybenzotriazole hydrate (#AS-65282) (Anaspec), *d*-Desthiobiotin (D1411) (MilliporeSigma), 4-Methyl Piperidine (#M73206) (MiliporeSigma), Rink Amide resin (#01-64-0013) (Novabiochem), Protected amino acids used in solid phase peptide synthesis (Novabiochem), Potassium Chloride (#P3911), Sodium Chloride (#S9888) (MilliporeSigma); 1M Tris-HCl buffer pH 7.5 (#15567027), Glycerol (#J61059), Imidazole (#A10221), Invitrogen Alexa FluorTM 594 hydrazide (#A10438), Tris (2-Carboxyethyl) Phosphine Hydrochloride (#50-153-2844) (Thermo Fisher Scientific), Zinc Chloride 1M Aqueous Solution (#470303-074) (Ward's Science).

Cell lines & plasmids

E. coli BL21(DE3) competent cells (#230132) and E. coli BL21-Gold(DE3) competent cells (#200131) (Agilent); all synthetic gBlocks (IDT, Table S1); pet-32a(+) vector (MilliporeSigma, #69015).

Cell culture/Microscopy

SsoFast™ EvaGreen® Supermix (#1725200, BioRad); McCoy's 5A Medium without phenol red (#SH30270, Cytiva); all siRNAs (Dharmacon, Table S1); RT-qPCR primers (IDT, Table S1); 4-well #1.5H glass bottom coverslip (#80427, Ibidi); Lipofectamine RNAiMAX transfection reagent (#13778150, Invitrogen); CellLight early endosomes, late endosomes, and lysosomes-GFP (BacMam 2.0; Life Technologies, catalog #C10586, #C10588, #C10596), Fibronectin bovine plasma (#F1141), Omni 0.5mL Tubes w/ 1.4mm Ceramic Beads (#19626) (MilliporeSigma); FuGENE HD transfection reagent (Promega, #E2311); GibcoTM Dulbecco's modified eagle medium (DMEM) without phenol red, high glucose, with 25 mM HEPES (#21063), Gibco™ Dulbecco's Modified Eagle Medium/Nutrient Mixture F-12 with (11320033) and without phenol red (21041025), Gibco™ GlutaMAXTM Supplement, 200 mM (#35050061), Gibco™ Sodium Pyruvate, 100 mM (#11360070), Gibco™ TrypLE Express Enzyme (1x) with (#12605010) and without (#12604013) phenol red, McCoy's 5a Medium (#16600082), Nunc™ Lab-Tek™ 8-well Chambered Coverglass (#155411) (Thermo Fisher Scientific); Saos-2 cell stock (UC Berkeley Cell Culture Facility), CHO/dhFr- cell stock (UC Berkeley Cell Culture Facility).

Molecular biology reagents

HiTrap® SP HP 5-mL column (#17115101), StrepTrap® HP 5-mL column (#28907547), PD-10 desalting columns (#17085101) (Cytiva); 3 kDa MWCO Amicon® Ultra-15 Centrifugal Filter Unit (#UFC90003), 10 kDa MWCO Amicon® Ultra-15 Centrifugal Filter Unit (#UFC9010), Benzonase® Nuclease HC (#71205), cOmplete, Mini EDTA-free protease inhibitor cocktail (#4693159001), ZipTips with 0.6 µL C4 resin (#ZTC04S008), 0.22 µm hydrophilic PVDF membrane filter (#SLGVR33RS, MilliporeSigma); NEBuilder® HiFi DNA Assembly Master Mix (#E2621L), Q5® High-Fidelity 2X Master Mix (#M0492S) (New England Biolabs); TALON® metal affinity resin (Takara, #635504); Pierce™ 660 nm Protein Assay Reagent (#22660), Pierce™ BCA Protein Assay Kit (#23227), Pierce™ Universal Nuclease for Cell Lysis (#88700) (Thermo Fisher Scientific), Dihydrofolate Reductase Assay Kit (CS0340) (Sigma Aldrich), CellTiter-Glo® 2.0 Assay Kit (#G9241) (Promega), His-tag antibody (Cell Signaling Technology, #2365), Tubulin antibody (CST, #2125), HRP-linked anti-rabbit IgG secondary antibody (CST, #7074).

#### II. Solid-phase peptide synthesis

##### A. Preparation of ZF5.3, CCHC, and AV5.3

AV5.3, CCHC, and ZF5.3 were synthesized using solid phase methods on a 0.1 mmol scale using a GPT PurePep Chorus automated peptide synthesizer. Rink Amide resin (100-200 mesh, Novabiochem) was swelled in DMF for 20 min at room temperature. Fmoc deprotection steps were performed twice using 5 mL of 20% 4-methyl piperidine in DMF at 50°C for 2 min.

Peptide coupling reactions were carried out by adding 5 equivalents of the necessary Fmoc-protected α-amino acid (Novabiochem), hexafluorophosphate benzotriazole tetramethyl uronium (Anaspec), 1-hydroxybenzotriazole hydrate (Anaspec), and 10 equivalents of DIPEA (Sigma-Aldrich) at 75°C for 7 min. This procedure was used for all amino acids except cysteine, histidine, and arginine, which were coupled at 50°C to minimize racemization, with arginine coupling repeated twice.

All peptides used in this paper carried amides at their C-termini and were acetylated at their N-termini in accord with previously synthesized sequences.[1,2] Peptides used for circular dichroism (CD) were cleaved from the resin and purified using reverse phase high-performance liquid chromatography (RP-HPLC) as previously described.[2] HPLC fractions containing the pure desired product were pooled and observed mass of the isolated material were examined using an Agilent 6530 QTOF LCMS. The pooled fractions were then lyophilized, and stored at -20˚C until use.

#### B. Preparation of ZF5.3$^{Rho}$ and AV5.3$^{Rho}$

AV5.3$^{Rho}$ and ZF5.3$^{Rho}$ were also synthesized using solid phase methods on a 0.1 mmol scale using a GPT PurePep Chorus automated peptide synthesizer and labeled on their N-termini with Lissamine rhodamine B as previously described.[3] Briefly, the sequences of the two peptides was extended at the N-terminus upon addition of a single (S)-2-(Fmoc-amino)-6-azidohexanoic acid residue and the terminal $NH_2$ group was acetylated to generate Ac-Lys($N_3$)-AV5.3 and Ac-Lys($N_3$)-ZF5.3. The peptides were cleaved from the resin for 4 h at RT using a mixture of 83% TFA, 5% phenol, 5% $H_2O$, 5% thioanisole, 1% DTT (w/v), and 1% TIPS.

After cleavage and purification by RP-HPLC, Ac-Lys($N_3$)-AV5.3 and Ac-Lys($N_3$)-ZF5.3 were reacted with Lissamine Rhodamine B functionalized with DBCO (Rho-DBCO). Rho-DBCO was synthesized as previously reported.[3] Briefly, AV5.3$^{Rho}$ and ZF5.3$^{Rho}$ were generated by reacting 1 equivalent of Ac-Lys($N_3$)-AV5.3 or Ac-Lys($N_3$)-AV5.3 with 10 equivalents of Rho-DBCO in DMSO for 1 h at RT. The products were purified by RP-HPLC, lyophilized, and stored at -20˚C until use.

#### C. Reconstitution of ZF5.3 and AV5.3 from stock solutions

All peptides were reconstituted from lyophilized powders in a Reconstitution Buffer containing 20 mM Tris-HCl, 150 mM KCl, pH 7.5. Cysteines were reduced for 10 min at RT by adding 1 molar equivalent of TCEP per cysteine residue. The peptides were subsequently coordinated with Zn(II) by adding two molar equivalents of $ZnCl_2$. Peptides used for circular dichroism were quantified by Pierce™ 660 Protein Assay. RhoB-labeled peptides were quantified using the Beer Lambert Law and the extinction coefficient for lissamine rhodamine B diethylamine that was determined to be 112,000 ± 2,000 cm-1 M-1 at λmax (569 nm).[2] Peptides were reconstituted when needed and used fresh to avoid peptide aggregation and precipitation. Peptides were routinely assessed for identity and purity using LC-MS.

### III. Circular dichroism

#### A. Instrumentation

Circular dichroism (CD) measurements were conducted and analyzed following established protocols.[4,5] Spectra were captured using a JASCO J-1500 Circular Dichroism Spectropolarimeter. A 0.1 cm pathlength quartz cuvette (Starna) was used for all recordings. The wavelength-dependent CD spectra were gathered from 300 to 200 nm at 1 nm intervals with a 5-sec averaging time, normalized to a baseline spectrum of the buffer alone.

#### B. pH-dependent assays

For samples where the  wavelength spectra was acquired across multiple pHs, the sample pH was adjusted with 0.1 M HCl or 0.1 M KOH. The pH measurements were done using a pH sensor InLab® Micro (Mettler-Toledo) before and after the spectra were recorded. Protein concentrations were also quantified after the spectra were recorded. All pH values were within ± 0.1 pH units.

##### C. Temperature-dependent assays

Temperature-dependent CD spectra were obtained from 5 to 90°C, with measurements taken every 2° and an equilibration time of 90 sec at each step. For wavelength spectra acquired across multiple pHs, the pH of the sample was adjusted with 0.1M HCl or 0.1M KOH and measured on a pH sensor InLab® Micro (Mettler-Toledo) before and after the spectra were recorded. All pH values were within ± 0.1 pH units. For wavelength spectra characterizing the Zn(II)-dependent fold, the peptides were reconstituted and the cysteine were reduced with TCEP, as described previously. Measurements were taken before and after the addition of 2 molar equivalents of Zn(II).

The wavelength for temperature melts was selected based on the wavelength spectra, set at 208 nm for AV5.3, ZF5.3, and CCHC, and 210 nm for DHFR proteins. AV5.3-DHFR and ZF5.3-DHFR spectra were collected at 20 μM protein concentration in a buffer of 25 mM Tris, 150 mM KCl, 1 mM TCEP, pH 7.5, 25 μM $ZnCl_2$. For experiments involving methotrexate (MTX), the proteins were pre-incubated with an equivalent amount of MTX for 30 min prior to measurement. Raw ellipticity values were converted to mean residue ellipticity using eq (1):

$$MRE\ [\theta] \ = \ \frac{m^{\circ} \cdot MRW}{l \cdot C} \qquad (1)$$

where θ = mean residue ellipticity in deg • $cm^2$ • $dmol^{-1}$, m° = raw ellipticity in millidegrees, MRW = mean residue weight (calculated as the molecular weight divided by the number of backbone amide bonds), $l$ = path length of the cuvette in millimeters, and C = concentration of the protein in mg $mL^{-1}$. For temperature melts, normalized intensity was calculated by normalizing all ellipticity values to the value at 5°C.

#### IV. Plasmids

##### A. Preparation of pAV-DHFR

pAV-DHFR was used to overexpress the AV5.3-DHFR fusion protein used in Figures (**4-5**). The gBlock was codon-optimized for expression in E. coli B strain and ordered from Integrated DNA Technologies (IDT). The gBlock for AV5.3-DHFR encoded the murine DHFR DNA sequence with an N-terminal AV5.3, separated by a glycine-serine-glycine (GSG) linker. Both sequences included a C-terminal LPETGG motif[5–8] for sortase conjugation and a $His_6$ tag for affinity purification. The gBlocks were inserted into a pET-32a(+) backbone vector (Sigma) linearized with Primers 1 and 2 using Gibson assembly. The identities of all plasmids were confirmed by Sanger sequencing and whole plasmid sequencing. Relevant DNA and protein sequences are listed in Table S1.

##### B. Preparation of pZF-DHFR

pZF-DHFR was used to overexpress the ZF5.3-DHFR fusion protein used in Figures (4-5). Plasmid generation was done as reported above. Relevant DNA and protein sequences are listed in Table S1.

#### V. Protein Expression, Purification, and Characterization

##### A. AV5.3-DHFR

Expression

The plasmid encoding AV5.3-DHFR were introduced into E. coli BL21(DE3) competent cells and grown on LB agar plates containing Carbenicillin (carb). A single colony from each plate was used to start a 20 mL overnight culture in LB media with 100 µg/mL carb, incubated at 37°C. The next morning, this starter culture was transferred to 1 L of LB with 100 mg/L amp and grown at 37°C until the optical density at 600 nm (OD600) reached 0.6-0.8. Protein expression was induced by adding 1 mM IPTG, and the cultures were incubated at 20°C for 20 hrs. Subsequent steps were carried out at 4°C.

Purification

The cells were harvested by centrifugation at 4000 g for 45 min and resuspended in 10 mL of ice-cold Lysis Buffer (25 mM Tris, 400 mM KCl, 5 mM TCEP, 5% glycerol, pH 7.5), supplemented with one cOmplete, mini EDTA-free protease inhibitor cocktail tablet and 4 µL of Benzonase HC. The cells were lysed using sonication for a total of 7 min, with 30-sec pulses followed by 30-sec rest intervals for 7 cycles. The lysate was clarified by centrifugation at 16,000 rpm for 40 min and then transferred to a 50 mL conical tube containing 2 mL of TALON® metal affinity resin pre-equilibrated with Wash Buffer 1 (25 mM Tris, 400 mM KCl, 15 mM imidazole, 5 mM TCEP, 5% glycerol, pH 7.5). The mixture was incubated with gentle rotation for 40 min.

After incubation, the lysate-resin mixture was transferred to a gravity column to drain the flowthrough. The resin was washed twice with 20 mL of Wash Buffer 2 (25 mM Tris, 400 mM KCl, 15 mM imidazole, 5 mM TCEP, 5% glycerol, pH 7.5) and twice with 20 mL of Wash Buffer 1. The protein was eluted with 14 mL of Elution Buffer (25 mM Tris, 300 mM KCl, 250 mM imidazole, 15% glycerol, pH 7.5), collected in 1 mL fractions. These fractions were analyzed by SDS-PAGE, and those containing pure protein were pooled and dialyzed overnight at 4°C in 1 L of Storage Buffer (25 mM Tris, 150 mM KCl, 1 mM TCEP, 15% glycerol, 100 µM $ZnCl_2$, pH 7.5). The following day, the protein concentration was determined using a Pierce™ BCA Protein Assay kit. If necessary, the protein was concentrated to over 1 mg/mL using an Amicon 10 kDa MWCO spin concentrator.

Characterization

The purified protein was characterized by using liquid chromatography-mass spectrometry (Agilent Infinity II LC/6530 Accurate-Mass Q-TOF LC/MS). The proteins were then stored at -80°C until use.

B. **ZF5.3-DHFR**

The expression, purification, and characterization of ZF5.3-DHFR was done by following directly a previously reported protocol for this protein.[5] Protein aliquots were stored at -80˚C.

**VI. Rhodamine labeling of proteins**

**A. AV5.3-DHFR$^{Rho}$**

Sortase reaction of AV5.3-DHFR and purification

AV5.3-DHFR$^{Rho}$ was produced through a reaction with a GGGK$^{Rho}$ peptide, utilizing the enzyme StrepTagII-SrtA7m-His$_6$. ZF5.3-DHFRRho was produced as previously described.[5] The GGGK$^{Rho}$ peptide synthesis followed a previously established method.[7]
For the labeling process of AV5.3-DHFR, the protein was diluted to 35 µM in a 1.5 mL solution containing 75 µM of the SrtA7m enzyme and 200 µM GGGK$^{Rho}$ in 20 mM Tris, 150 mM KCl, 1 mM TCEP, pH 7.5. The reaction was gently rotated for 4 h at 4°C. Post-reaction, AV5.3-DHFR$^{Rho}$ was isolated by incubating with 1 mL TALON® resin for 1 h at 4°C, followed by application to a gravity column. The column was washed with 10 mL of Wash Buffer 2. Since the His6 tag is located C-terminal to the LPETGG motif on DHFR, a successful sortase reaction results in the loss of the His$_6$ tag, and thus, AV5.3-DHFR$^{Rho}$ is found in the column washes and flowthrough. The fractions containing fluorescent AV5.3-DHFR were analyzed via SDS-PAGE using a fluorescence imager, pooled, and dialyzed to 25 mM Tris, 150 mM KCl, 1 mM TCEP, 15% glycerol, pH 7.5. To further purify AV5.3-DHFR$^{Rho}$ it was run through a StrepTrap XP column (Cytiva) to ensure SrtA7m was removed, The flow though was collected and buffer exchanged to Storage Buffer.

Characterization of AV5.3-DHFR$^{Rho}$

The purified AV5.3-DHFR$^{Rho}$ was characterized by LC/MS. Protein concentrations were measured using the Pierce™ 660 nm Protein Assay (Sigma). The concentration of rhodamine in labeled protein was determined using the Beer-Lambert Law, where the absorbance was measured at 570 nm on a NanoDrop spectrophotometer. The extinction coefficient of lissamine rhodamine B in water is 112000 $M^{-1}$ $cm^{-1}$. To calculate labeling efficiency the rhodamine concentration was divided by the total protein concentration. Protein was stored at -80°C.

##### B. ZF5.3-DHFR$^{Rho}$

The rhodamine labeling of ZF5.3-DHFR was performed by following a protocol reported previously.[5] The material was characterized as described above. Protein was stored at -80˚C.

#### VII. Delivery methods

##### A. Cell culture

Cell stocks of Saos-2 were purchased from the UC Berkeley Cell Culture Facility. Saos-2 cells were cultured in complete media composed of McCoy's 5A medium with phenol red containing 15% fetal bovine serum (FBS), penicillin and streptomycin (P/S, 100 units/mL and 100 µg/mL, respectively), 1 mM sodium pyruvate, and 2 mM GlutaMax.

##### B. Delivery procedure

All fluorescence-based intracellular delivery experiments, including confocal microscopy, flow cytometry, and fluorescence correlation spectroscopy, were conducted following established protocols.[9] One day prior to the experiments, 2.0 x $10^5$ Saos-2 cells were seeded in a 6-well dish with Clear McCoy's 5A medium (no phenol red) containing 15% FBS. The next day, cells were treated with 1 mL of ZF5.3, AV5.3, or DHFR proteins, diluted to a concentration of 0.1 - 1 µM in clear McCoy's 5A medium. Concurrently, a 0.1 mg/mL fibronectin solution in DPBS was added to designated wells of an 8-well microscopy dish. To determine the microscope's focal volume for FCS, 400 µL of a 100 nM AlexaFluor 594 hydrazide dye solution in water was added to another well. The dishes were maintained at 37°C with 5% $CO_2$ during cell incubation and subsequent steps. Five min before the incubation ended (lasting 0.5-1 hr), Hoechst 33342 nuclear dye was added to each well of the 6-well dish at a final concentration of 300 nM. After incubation, cells were washed three times with 2 mL DPBS. Cells were trypsinized with 500 µL trypsin (without phenol red). The trypsinization was quenched using 1 mL of clear McCoy's 5A medium + 15% FBS to the well. The samples were collected in 15 mL Falcon tubes and the well were washed with another 1.0 mL of clear McCoy's 5A medium + 15% FBS. Cells were then pelleted and washed once with 1 mL clear DMEM + Hepes. During this centrifugation step, the microscopy dish was removed from the incubator, and the fibronectin-coated wells were washed three times with 200 µL DPBS, leaving DPBS in the wells until cell addition. The pelleted cells were resuspended in 600 µL DMEM + Hepes, with 300 µL added to each fibronectin-coated well for confocal microscopy and FCS analysis, then incubated at 37°C with 5% $CO_2$ to allow cells to adhere. The remaining 300 µL of cells were pelleted, resuspended in 1 mL DPBS, and transferred to a 1.5 mL Eppendorf tube for flow cytometry.

##### C. Confocal microscopy

General procedures for confocal microscopy have been previously detailed.[8,9]

Colocalization Experiment

Saos-2 cells were seeded into tissue culture-treated 6-well plates at a density of 100,000 cells per well in complete McCoy's 5A medium without phenol red (2 mL/well, supplemented with 15% FBS and 1 mM sodium pyruvate, but without penicillin/streptomycin) 48 hours prior to the experiment. The next day, the cells were transduced with CellLight early endosomes, late endosomes, or lysosomes-GFP following the manufacturer's instructions. This transduction was incubated for 18 hours. After incubation, the media were aspirated, and the cells were washed twice with DPBS. Subsequently, the cells were incubated with 1μM AV5.3 in clear McCoy's 5A medium without phenol red for 30 minutes at 37 °C in a 5% $CO_2$ atmosphere. During this incubation, an 8-well microscope slide was prepared for microscopy as described in **Delivery Procedure**. After the 30-minute AV5.3 incubation, the cells' nuclei were stained with Hoechst 33342 (300 nM) for 5 minutes at 37 °C in 5% $CO_2$. Cells were then washed and plated for microscopy, also thoroughly described in **Delivery Procedure**. Imaging was performed using a Leica DMi8 CS scanhead with the following parameters: Hoechst excitation at 405 nm (emission 410–496 nm), eGFP excitation at 488 nm (emission 490–526 nm), and Lissamine RhoB excitation at 561 nm (emission 570–607 nm). The degree of colocalization between two fluorescence channels was quantified using the Pearson's r correlation coefficient in FIJI software.

##### D. Flow cytometry

For each sample, 20,000 cells were analyzed via flow cytometry following the described protocol. Forward and side scatter gating parameters were used to select whole cells, based on previously established settings.[5,7,8] The fluorescence intensities for Hoechst 33342 (excited with a 405 nm violet laser, emission detected at 440 ± 50 nm) and lissamine rhodamine B (excited with a 561 nm yellow laser, emission detected at 586 ± 16 nm) were measured using an Attune NxT Focusing Flow Cytometer. The median fluorescence intensity (MFI) for the lissamine rhodamine B channel was recorded for the gated cell population and adjusted for the rhodamine labeling efficiency if needed. Each flow cytometry experiment included at least two biological replicates.

##### E. Fluorescence correlation spectroscopy (FCS)

The procedures for confocal microscopy and fluorescence correlation spectroscopy (FCS) have been previously detailed.[8,9] Experiments were conducted using a STELLARIS 8 microscope (Leica Microsystems) equipped with a Leica DMi8 CS scanhead. A HC Plan-Apo 63x/1.4NA water immersion objective was used for FCS measurements. The system included a pulsed white-light laser (440 nm to 790 nm; power: 440 nm > 1.1 mW, 488 nm > 1.6 mW, 560 nm > 2.0 mW, 630 nm > 2.6 mW, 790 nm > 3.5 mW, 78 MHz) and a pulsed 775 nm STED laser. Confocal imaging utilized HyD S or HyD X detectors in analog mode, while FCS measurements

exclusively used the Hybrid HyD X detector. All microscopy experiments were performed at 37°C with 5% $CO_2$ in a blackout cage enclosure (Okolab), monitored by Oko-Touch.
Before each experiment, the objective's correction collar was adjusted to maximize the counts per molecule for the AlexaFluor 594 hydrazide dye standard, accounting for minor fluctuations due to varying glass-bottom LabTek™ dish thicknesses. Lissamine rhodamine B was excited at 561 nm with a fluorescence filter of 570 - 660 nm, and the laser's pinhole S5 was set to 1 AU. Prior and after in-cell experiments, ten five-sec autocorrelation traces were obtained from a well containing 100 nM AlexaFluor 594 hydrazide dye to calculate the microscope's focal volume.

For in-cell data collection, confocal microscopy images were used to position the microscope laser's crosshairs in the cytosol of the cells, avoiding areas with punctate fluorescence indicative of endosomal entrapment. Each FCS measurement consisted of ten five-sec traces, with at least 10 cells per condition measured for each biological replicate, and a minimum of two biological replicates per condition. Expected diffusion times for rhodamine-labeled ZF5.3, AV5.3 and DHFR proteins were determined by measuring in vitro autocorrelation traces of 50-200 nM protein solutions in clear DMEM media (+25 mM Hepes) at 37°C.

##### F. FCS data analysis

Autocorrelation traces obtained from fluorescence correlation spectroscopy (FCS) measurements were analyzed using a custom MATLAB script.[6–9] To extract quantitative information from *in cellula* data, it was essential to determine the effective confocal volume of the microscope. This value was established using equations 4-6 and the in vitro autocorrelation traces for the AlexaFluor 594 hydrazide standard, which has a known diffusion coefficient in water[2] and was measured at the start of each experiment. These traces were fitted to a 3D diffusion equation (2):

$$G(\tau) = \frac{1}{N} \cdot \frac{1}{(1+\frac{\tau}{\tau_{diff}})\sqrt{1+(s^2\frac{\tau}{\tau_{diff}})}} \qquad (2)$$

where N = the average number of molecules detected in the focal volume ($V_{eff}$), $\tau_{diff}$ = the average diffusion time that a molecule requires to cross $V_{eff}$, and s = the structure factor (the ratio of the radial to axial dimensions of the focal volume). The structure factor was measured to be 0.17 using the autocorrelation function of AlexaFluor 594 in water at 25°C and fixed for all subsequent analyses. $V_{eff}$ can be extracted from these data by inserting the $\tau_{diff}$ value derived from eq (2) to calculate $\omega_1$ in eq (3):

$$\omega_1 = \sqrt{4 \cdot D \cdot \tau_{diff}} \qquad (3)$$

where $\omega_1$ = the lateral extension of the confocal volume and D = the known diffusion coefficient of AlexaFluor 594 in water at 37°C ($5.20 \times 10^{-6}$ $cm^2$ $s^{-1}$). $V_{eff}$ can then be directly calculated from eq (4):

$$V_{eff} = \pi^{\frac{3}{2}} \cdot \omega_1^{3} \cdot \frac{1}{s} \qquad (4)$$

The average $V_{eff}$ for all experiments ranged from 0.25 to 0.4 fL. Autocorrelation traces derived from in-cellulo measurements were fitted using a 3D anomalous diffusion equation (5):

$$G(\tau) = \frac{1}{N} \cdot \frac{1}{(1+\frac{\tau}{\tau_{diff}})^{\alpha}\sqrt{1+(s^2\frac{\tau}{\tau_{diff}})^{\alpha}}} + G(\infty) \qquad (5)$$

where N = the average number of molecules in the focal volume, $\tau_{diff}$ = the average diffusion time that a molecule requires to cross $V_{eff}$, $\alpha$ = the anomalous diffusion coefficient, and s = the structure factor (0.17). Parameters resulting from the fitting, including diffusion time, anomalous diffusion coefficient, and counts per molecule, were filtered as previously reported, and only fitted curves that passed this filtering and displayed Chi-squared values under 100 were selected for further analysis. Expected $\tau_{diff}$ values for *in cellula* samples were predicted from autocorrelation traces for in vitro protein samples in DMEM that were fitted using equation 3. Only curves that displayed in-cellulo $\tau_{diff}$ values within 2-20-fold of the in vitro value were selected. Shorter diffusion times can indicate degradation, while longer diffusion times may reflect aggregated material with abnormal diffusion dynamics. The concentration (CC) of protein in the cytosol was then calculated using the value of N derived above in equation 6:

$$C = \frac{N}{N_A \cdot V_{eff}} \qquad (6)$$

where $N_A$ = Avogadro's number ($6.023 \times 10^{23}$ $mol^{-1}$). When relevant, the concentration values were adjusted for fluorophore labeling efficiency. At least 10 concentration values from curves that passed all filters were used for each FCS condition (Table S2).

#### VIII. Endosomal leakage assay

One day prior to the experiment, Saos-2 cells were seeded into tissue culture-treated 6-well plates at a density of 100,000 cells per well in complete McCoy's 5A medium without phenol red (2 mL/well, supplemented with 15% FBS and 1 mM sodium pyruvate, but without penicillin/streptomycin). On the day of the experiment, the medium was removed, and cells were washed twice with DPBS. The cells were incubated with $Lys9^{Rho}$ (600 nM) in clear McCoy's 5A medium without phenol red (1 mL/well, without FBS, sodium pyruvate, or penicillin/streptomycin) for 30 minutes at 37°C in a 5% $CO_2$ atmosphere. During this incubation, an 8-well microscope slide was prepared as described in **Delivery Procedure**. Nuclei were labeled with Hoechst 33342 (300 nM) for 5 minutes at the end of the 1-hour incubation by adding the nuclear stain directly to the reagent-containing medium. Cells were then prepared for microscopy as described in **Delivery Procedure**. Lastly, the cells were allowed to adhere for 20 minutes at 37 °C in 5% $CO_2$ before FCS measurements were collected as described above.

#### IX. siRNA knockdowns

##### A. Knockdown of Saos-2 cells for delivery experiments

Knockdowns for HOPS and CORVET were conducted using the siRNAs specified in Table Sxx, following previously established methods.[2] Four days prior to delivery, 8.5 x $10^4$ Saos-2 cells were seeded in a 6-well dish with clear McCoy's 5A medium supplemented with 15% FBS (no P/S). The next day, cells were transfected with the target siRNA at a final concentration of 100 nM, using Lipofectamine RNAi/MAX in clear McCoy's 5A medium with 15% FBS (no P/S), and incubated for 4 h at 37°C with 5% $CO_2$. Following transfection, cells were washed three times with DPBS and incubated for 72 h at 37°C, 5% $CO_2$ in 2 mL clear McCoy's 5A medium with 15% FBS (no P/S) per well before starting the delivery experiments. To verify siRNA knockdown after 72 h, RT-qPCR analysis was conducted.[2,5]

##### B. siRNA knockdown validation by qPCR

The cDNA from Saos-2 cells was amplified using gene-specific primers (150 nM, see Table S1) with SsoFast EvaGreen Supermix (Bio-Rad) on a Bio-Rad CFX96 RT-PCR detection system. The experiment included three biological replicates for each knockdown, with each sample analyzed in triplicate. Each RT-qPCR analysis featured the target gene, GAPDH as a reference gene, and negative controls lacking either reverse transcriptase or template cDNA.

#### X. DHFR activity assays

##### A. *In vitro* sample preparation

To quantify the activity of wildtype DHFR (hDHFR), we used a DHFR aliquot purchased from Sigma. The activity of this hDHFR was used as a control and standard for the quantified activity of AV5.3-DHFR and ZF5.3-DHFR. Purified AV5.3-DHFR or ZF5.3-DHFR was accurately quantified using Pierce 660 and then added to a final concentration of 100 nM in a 200 µL solution of 1X activity assay buffer (Sigma).

##### B. Cytosolic fractionation method

The day before the experiment, 5.0×$10^6$ Saos-2 cells were seeded in a 150 mm dish containing clear McCoy's 5A medium with 15% FBS (no P/S). The following morning, cells were washed three times with DPBS. The medium was then replaced with 10 mL of clear McCoy's 5A medium (without FBS or P/S) containing a 1 µM solution of eitherAV5.3-DHFR or ZF5.3-DHFR. The cells were incubated at 37°C with 5% $CO_2$ for 1 hr.

Cell harvesting

After the incubation period, the cells were washed three times with DPBS and detached from the dish using trypsin. The trypsin reaction was stopped by adding 15 mL of clear McCoy's 5A medium supplemented with 15% FBS (no P/S). The cells were then transferred to a 50 mL falcon tube, and the dish was rinsed with an additional 5 mL of the same medium to collect any remaining cells. The cells were pelleted by centrifugation at 200 g for 3 min, then washed by resuspension in 3 mL of DPBS and pelleted again.

Cell lysis and fractionation

The cell pellet was resuspended in 1 mL of precooled buffered isotonic sucrose (290 mM sucrose, 10 mM imidazole pH 7.0, 1 mM DTT, and 1 cOmplete protease inhibitor tablet) and pelleted again. The cells were then resuspended in 100 µL of the isotonic sucrose buffer and transferred to 0.5 mL tubes containing 1.4 mm ceramic beads (Omni International). Using a Bead Ruptor, the cells were lysed for 10 sec at speed 1. The homogenized cells were transferred to polycarbonate ultracentrifuge tubes and the cytosolic fraction was separated from the membrane fraction by centrifugation at 350,000 g for 1 hr at 4°C (using a Beckman Coulter, TLA-100 rotor).

Western Blot

Samples for Western blot analysis were prepared by combining 20 µL of protein standards or cytosolic fractions with 5 µL of 5X SDS gel loading dye, followed by boiling at 95°C for 5 minutes. Western blotting was conducted using primary antibodies against His-tag (Cell Signaling Technology, #2365) and tubulin (CST, #2125). Detection was performed with an HRP-linked anti-rabbit IgG secondary antibody (CST, #7074).

Cytosolic fraction sample preparation for activity assay

The supernatant (cytosolic fraction) was immediately removed and transferred to a 1.5 mL Eppendorf tube. The total protein concentration was determined using Pierce™ 660nm protein assay kit (Sigma Aldrich). Cytosolic fraction solutions of 0.250 mg/mL were prepared for activity assay.

##### C. DHFR activity assay

The catalytic activities of AV5.3-DHFR and ZF5.3-DHFR were assessed using a commercially available Dihydrofolate Reductase Assay kit (Sigma). The assay monitors the NADPH-dependent reduction of dihydrofolate (DHFA) to tetrahydrofolate (THFA) by measuring the loss in absorbance at 340 nm over time using a UV/Vis Microplate Reader (Agilent BioTek Synergy HTX Multi-Mode Microplate Reader). Purified AV5.3-DHFR or ZF5.3-DHFR was added to a final concentration of 100 nM in a 200 µL solution of 1x activity assay buffer provided in the kit. The solution was added in the microplate, followed by addition of 1.2 µL of 10 mM NADPH. The plate was shaken to mix the reaction components. In quick succession, 1 µL of 10 mM DHFA was added, the plate was shaken again, and the reaction progress was monitored by absorbance at 340 nm. Absorbance readings were taken every 15 sec for 150 sec. For measurements in the presence of MTX, MTX was added to a final concentration of 100 nM and incubated with the protein for 5 min before addition of NADPH and DHFA. The activity of cytosolic fraction (CF)  samples was quantified by diluting 10 µL of CF to a final 200 µL reaction volume. The specific activity (in units per mg) of each sample was calculated using eq (7):

$$Units/mg = \frac{(\Delta OD/min(sample) - \Delta OD/min(blank)) \cdot d}{12.3 \cdot V \cdot mg/mL} \qquad (7)$$

where ΔOD/min(sample) = the slope of the 340 nm absorbance curve for the sample, ΔOD/min(blank) = the slope of the 340 nm absorbance curve for a buffer blank, d = dilution factor of the enzyme sample, 12.3 = the extinction coefficient ($mM^{-1}$ $cm^{-1}$) for the DHFR reaction at 340 nm, V = the volume of enzyme used in the reaction, and mg/mL = protein concentration in mg $mL^{-1}$.

##### D. DHFR rescue assay

To test if DHFR rescue in CHO/dhFr- cells would improve cell viability, we made use of the CellTiter-Glo® 2.0 cell viability assay (Promega). It detects viable cells by measuring ATP activity. In a fibronectin-treated 96-well plate we seeded $5.0 \times 10^3$ cells per condition replicate in clear IMDM media + 10% FBS + 1X (100 µM hypoxanthine/16 µM thymidine) HT supplement. Cells were let to adhere for 4 hrs. The media was then replaced with clear IMDM media + 10% FBS including varying HT supplement dosages tested. Then, 1 µM AV5.3-DHFR or 1µM ZF5.3-DHFR was treated for 1 hr in clear IMDM media. After 12 hrs, we took our first time point measurement (0 hrs) by using a UV/Vis Microplate Reader (Agilent BioTek Synergy HTX Multi-Mode Microplate Reader). We re-dosed DHFR proteins every 12 hrs for 1 hr. Results were represented in cell viability percentage where the control was cells treated with just 1X HT supplement.

#### XI. Statistics

All statistical tests were performed with the GraphPad Prism 9 software. All details are described in their appropriate figure legends. Flow cytometry and FCS data were analyzed using Brown-Forsythe and Welch ANOVA statistical tests followed by unpaired t-tests with Welch’s correction.

**XII. Supplementary Tables 1-3**

**Table S1:** Relevant Peptide, Protein, and DNA/RNA Sequences

| | | |
|---|---|---|
| **Solid-Phase Peptide Sequences** | | |
| ZF5.3 | 3422 Da | Ac-YSCNVCGKAFVLSRHLNRHLRVHRRAT-$NH_2$ |
| ZF5.3$^{Rho}$ | 4277 Da | Ac-K$^{Rho}$-YSCNVCGKAFVLSRHLNRHLRVHRRAT-$NH_2$ |
| AV5.3 | 3354 Da | Ac-YSCNVCGKAFVLSRHLNRCLRVCRRAT-$NH_2$ |
| AV5.3$^{Rho}$ | 4210 Da | Ac-K$^{Rho}$-YSCNVCGKAFVLSRHLNRCLRVCRRAT-$NH_2$ |
| CCHC | 3388 Da | Ac-YSCNVCGKAFVLSRHLNRHLRVCRRAT-$NH_2$ |
| **Protein Sequences** | | |
| ZF5.3-DHFR | 26,246 Da | YSCNVCGKAFVLSRHLNRHLRVHRRATGSGVRPLNCIVAVS QNMGIGKNGDLPWPPLRNEFKYFQRMTTTSSVEGKQNLVI MGRKTWFSIPEKNRPLKDRINIVLSRELKEPPRGAHFLAKSL DDALRLIEQPELASKVDMVWIVGGSSVYQEAMNQPGHLRL FVTRIMQEFESDTFFPEIDLGKYKLLPEYPGVLSEVQEEKGI KYKFEVYEKKLPETGGHHHHHH |
| ZF5.3-DHFR$^{Rho}$ | 26,147 Da | YSCNVCGKAFVLSRHLNRHLRVHRRATGSGVRPLNCIVAVS QNMGIGKNGDLPWPPLRNEFKYFQRMTTTSSVEGKQNLVI MGRKTWFSIPEKNRPLKDRINIVLSRELKEPPRGAHFLAKSL DDALRLIEQPELASKVDMVWIVGGSSVYQEAMNQPGHLRL FVTRIMQEFESDTFFPEIDLGKYKLLPEYPGVLSEVQEEKGI KYKFEVYEKKLPETG-GGGK$^{Rho}$ |
| AV5.3-DHFR | 26,178 Da | YSCNVCGKAFVLSRHLNRCLRVCRRATGSGVRPLNCIVAVS QNMGIGKNGDLPWPPLRNEFKYFQRMTTTSSVEGKQNLVI MGRKTWFSIPEKNRPLKDRINIVLSRELKEPPRGAHFLAKSL DDALRLIEQPELASKVDMVWIVGGSSVYQEAMNQPGHLRL FVTRIMQEFESDTFFPEIDLGKYKLLPEYPGVLSEVQEEKGI KYKFEVYEKKLPETGGHHHHHH |
| AV5.3-DHFR$^{Rho}$ | 26,080 Da | YSCNVCGKAFVLSRHLNRCLRVCRRATGSGVRPLNCIVAVS QNMGIGKNGDLPWPPLRNEFKYFQRMTTTSSVEGKQNLVI MGRKTWFSIPEKNRPLKDRINIVLSRELKEPPRGAHFLAKSL DDALRLIEQPELASKVDMVWIVGGSSVYQEAMNQPGHLRL FVTRIMQEFESDTFFPEIDLGKYKLLPEYPGVLSEVQEEKGI KYKFEVYEKKLPETG-GGGK$^{Rho}$ |
| **Primers** | | |
| Primer 1 (to linearize pet-32a(+) vector) | | ATGTATATCTCCTTCTTAAAGTTAAACAAAATTATT |

| | |
|---|---|
| Primer 2 (to linearize pet-32a(+) vector) | TAACAAAGCCCGAAAGGAAG |
| **siRNA used for RNAi** | |
| RISC-free | siGENOME RISC-Free Control siRNA: D-001220-01 (Dharmacon) |
| VPS39 | siGENOME human SMARTpool siRNA VPS39: M-014052-01 (Dharmacon) |
| VPS41 | siGENOME human SMARTpool siRNA VPS41:M-006972-01 (Dharmacon) |
| TGF-BRAP1 | siGENOME human SMARTpool siRNA TGFBRAP1: M-006903-01 (Dharmacon) |
| VPS8 | siGENOME human SMARTpool siRNA VPS8: M-023668-02 (Dharmacon) |
| **Primers for RT-qPCR** | |
| GAPDH | PrimeTime qPCR primers Hs.PT.39a.22214836 (IDT) |
| VPS39 | PrimeTime qPCR primers Hs.PT.58.26949378 (IDT) |
| VPS41 | PrimeTime qPCR primers Hs.PT.58.39633303 (IDT) |
| VPS8 | PrimeTime qPCR primers Hs.PT.58.1611351 (IDT) |
| TGF-BRAP1 (forward) | CTGACCACTCAGTACATCATCC |
| TGF-BRAP1 (reverse) | CTCCTGTCTCCCTATCCTCTT |

**Table S2:** Summary Values for Intracellular Delivery Experiments

*In flow cytometry (FC) experiments, n refers to the number of biological replicates. For each biological replicate, 20,000 cells were measured. The "FC Mean" corresponds to the average Median Fluorescence Intensity recorded across all biological replicates. For fluorescence correlation spectroscopy (FCS), n refers to the number of individual cell measurements that passed all final filtering criteria (see Methods for more details) across all biological replicates. The number of biological replicates for FCS experiments is the same as that shown for the corresponding flow cytometry experiment. The "FCS Mean" corresponds to the average cytosolic concentration measured for that condition.*

| Condition | Fig. | n - FC | FC Mean | FC SEM | n - FCS | FCS Mean | FCS SEM |
|---|---|---|---|---|---|---|---|
| ZF5.3$^{Rho}$ - 0.1 µM, 30 min | 2 | 2 | 1023 | 30.00 | 43 | 68.70 | 7.872 |
| ZF5.3$^{Rho}$ - 0.5 µM, 30 min | | 2 | 5854 | 3.500 | 12 | 259.9 | 48.67 |
| ZF5.3$^{Rho}$ - 1.0 µM, 30 min | | 2 | 10299 | 98.50 | 15 | 405.7 | 42.85 |
| AV5.3$^{Rho}$ - 0.1 µM, 30 min | | 3 | 493.3 | 22.00 | 57 | 58.75 | 4.942 |
| AV5.3$^{Rho}$ - 0.5 µM, 30 min | | 3 | 1627 | 132.4 | 37 | 201.3 | 21.20 |
| AV5.3$^{Rho}$ - 1.0 µM, 30 min | | 3 | 3228 | 274.1 | 26 | 439.2 | 58.46 |
| ZF5.3$^{Rho}$ - RISC-free siRNA, 1 µM, 30 min | 3 | 4 | 12252 | 1536 | 26 | 454.7 | 33.04 |
| ZF5.3$^{Rho}$ - VPS8 siRNA, 1 µM, 30 min | | 2 | 9234 | 144.5 | 22 | 584.0 | 30.55 |
| ZF5.3$^{Rho}$ - TGF-BRAP1 siRNA, 1 µM, 30 min | | 2 | 22127 | 292.5 | 23 | 717.7 | 62.33 |
| ZF5.3$^{Rho}$ - VPS39 siRNA, 1 µM, 30 min | | 2 | 9274 | 294.0 | 21 | 156.7 | 14.97 |
| ZF5.3$^{Rho}$ - VPS41 siRNA, 1 µM, 30 min | | 2 | 7101 | 365.5 | 20 | 203.2 | 21.45 |
| AV5.3$^{Rho}$ - RISC-free siRNA, 1 µM, 30 min | | 4 | 3006 | 129.1 | 50 | 298.9 | 29.57 |
| AV5.3$^{Rho}$ - Lipofectamine control, 1 µM, 30 min | | 2 | 2833 | 79.50 | — | — | — |
| AV5.3$^{Rho}$ - VPS8 siRNA, 1 µM, 30 min | | 2 | 572.0 | 36.00 | 30 | 78.89 | 9.345 |
| AV5.3$^{Rho}$ - TGF-BRAP1 siRNA, 1 µM, 30 min | | 2 | 1422 | 9.000 | 19 | 23.97 | 2.821 |

| | | | | | | | |
|---|---|---|---|---|---|---|---|
| AV5.3$^{Rho}$ - VPS39 siRNA, 1 µM, 30 min | | 2 | 1106 | 2.000 | 34 | 42.10 | 3.852 |
| AV5.3$^{Rho}$ - VPS41 siRNA, 1 µM, 30 min | | 2 | 451.5 | 14.50 | 15 | 38.64 | 6.580 |
| ZF5.3-DHFR$^{Rho}$ - 0.1 µM, 1 hr | 4 | 2 | 1583 | 123.5 | 30 | 107.3 | 10.64 |
| ZF5.3-DHFR$^{Rho}$ - 0.5 µM, 1 hr | | 2 | 10430 | 76.50 | 34 | 239.7 | 27.00 |
| ZF5.3-DHFR$^{Rho}$ - 1.0 µM, 1 hr | | 2 | 23389 | 1836 | 36 | 291.2 | 22.10 |
| AV5.3-DHFR$^{Rho}$ - 0.1 µM, 1 hr | | 3 | 1919 | 519.9 | 62 | 71.68 | 4.732 |
| AV5.3-DHFR$^{Rho}$ - 0.5 µM, 1 hr | | 3 | 13565 | 820.7 | 96 | 236.8 | 11.53 |
| AV5.3-DHFR$^{Rho}$ - 1.0 µM, 1 hr | | 3 | 28322 | 3632 | 50 | 329.1 | 36.11 |
| AV5.3-DHFR$^{Rho}$ - RISC-free siRNA, 1 µM, 1 hr | | 2 | 18885 | 2267 | 58 | 193.8 | 20.03 |
| AV5.3-DHFR$^{Rho}$ - Lipofectamine control, 1 µM, 1 hr | | 2 | 18461 | 4000 | — | — | — |
| AV5.3-DHFR$^{Rho}$ - VPS39 siRNA, 1 µM, 1 hr | | 2 | 16128 | 1385 | 15 | 104.9 | 11.62 |
| AV5.3-DHFR$^{Rho}$ - TGF-BRAP1 siRNA, 1 µM, 1 hr | | 2 | 20086 | 1862 | 42 | 101.2 | 5.109 |
| Lys9$^{Rho}$, 0.6 µM, 1.5 hr | S4 | – | – | – | 27 | 6.309 | 0.4880 |
| Lys9$^{Rho}$, 0.6 µM, 0.5 hr, + 1 µM ZF5.3, 1 hr | | – | – | – | 26 | 7.952 | 0.7563 |
| Lys9$^{Rho}$, 0.6 µM, 0.5 hr, + 1 µM AV5.3, 1 hr | | – | – | – | 28 | 8.624 | 0.6530 |
| Lys9$^{Rho}$, 0.6 µM, 0.5 hr + 16 µL/mL RNAiMAX, 1 hr | | – | – | – | 39 | 51.90 | 6.645 |
| AV5.3-DHFR$^{Rho}$ - 0.1 µM, 1 hr + MTX | S7 | 4 | 1036 | 219.8 | 50 | 66.32 | 4.500 |
| AV5.3-DHFR$^{Rho}$ - 0.5 µM, 1 hr + MTX | | 4 | 6374 | 554.2 | 95 | 113.9 | 7.563 |
| AV5.3-DHFR$^{Rho}$ - 1.0 µM, 1 hr + MTX | | 3 | 18296 | 527.6 | 52 | 148.9 | 11.63 |

**Table S3:** Statistical information for all comparisons

*All statistical tests presented were performed with the use of GraphPad Prism 9 software. All FCS data were analyzed using Brown-Forsythe and Welch ANOVA statistical tests followed by unpaired t-tests with Welch's correction. All relevant parameters are described below.*

| Comparison | Fig. | Mean Difference | 95.00% CI | p-value | Test statistic (t) | Degrees of Freedom |
|---|---|---|---|---|---|---|
| ZF5.3$^{Rho}$ vs AV5.3$^{Rho}$ - 0.1 µM, 30 min | 2B | 529.7 | 374.4 to 684.9 | 0.0043 | 14.24 | 2.066 |
| ZF5.3$^{Rho}$ vs AV5.3$^{Rho}$ - 0.5 µM, 30 min | | 4227 | 3657 to 4796 | 0.0010 | 31.91 | 2.003 |
| ZF5.3$^{Rho}$ vs AV5.3$^{Rho}$ - 1.0 µM, 30 min | | 7070 | 6019 to 8121 | 0.0005 | 24.27 | 2.468 |
| ZF5.3$^{Rho}$ vs AV5.3$^{Rho}$ - 0.1 µM, 30 min | 2C | 9.953 | -8.570 to 28.48 | 0.2878 | 1.071 | 73.11 |
| ZF5.3$^{Rho}$ vs AV5.3$^{Rho}$ - 0.5 µM, 30 min | | 58.67 | -54.22 to 171.6 | 0.2860 | 1.105 | 15.40 |
| ZF5.3$^{Rho}$ vs AV5.3$^{Rho}$ - 1.0 µM, 30 min | | -33.52 | -180.1 to 113.1 | 0.6463 | 0.4625 | 38.98 |
| ZF5.3$^{Rho}$ - RISC-free siRNA vs VPS8 siRNA, 1 µM, 30 min | 3B | -3018 | -7881 to 1844 | 0.1438 | 1.956 | 3.053 |
| ZF5.3$^{Rho}$ - RISC-free siRNA vs TGF-BRAP1 siRNA, 1 µM, 30 min | | 9875 | 5077 to 14673 | 0.0065 | 6.315 | 3.209 |
| ZF5.3$^{Rho}$ - RISC-free siRNA vs VPS39 siRNA, 1 µM, 30 min | | -2978 | -7775 to 1819 | 0.1470 | 1.904 | 3.211 |
| ZF5.3$^{Rho}$ - RISC-free siRNA vs VPS41 siRNA, 1 µM, 30 min | | -5151 | -9915 to -387.7 | 0.0407 | 3.262 | 3.317 |
| ZF5.3$^{Rho}$ - RISC-free siRNA vs VPS8 siRNA, 1 µM, 30 min | 3C | 129.4 | 38.82 to 220.0 | 0.0061 | 2.876 | 46.00 |
| ZF5.3$^{Rho}$ - RISC-free siRNA vs TGF-BRAP1 siRNA, 1 µM, 30 min | | 263.1 | 119.7 to 406.5 | 0.0007 | 3.729 | 33.75 |
| ZF5.3$^{Rho}$ - RISC-free | | -371.6 | -371.6 to | <0.0001 | 8.214 | 34.51 |

| | | | | | | |
|---|---|---|---|---|---|---|
| siRNA vs VPS39 siRNA, 1 µM, 30 min | | | -224.2 | | | |
| ZF5.3$^{Rho}$ - RISC-free siRNA vs VPS41 siRNA, 1 µM, 30 min | | -251.5 | -331.0 to -171.9 | <0.0001 | 6.385 | 40.95 |
| AV5.3$^{Rho}$ - RISC-free siRNA vs Lipofectamine control, 1 µM, 30 min | 3E | -173.5 | -595.1 to 248.1 | 0.3166 | 1.144 | 3.987 |
| AV5.3$^{Rho}$ - VPS8 siRNA vs TGF-BRAP1 siRNA, 1 µM, 30 min | | -850.0 | -1214 to -485.6 | 0.0194 | 22.91 | 1.125 |
| AV5.3$^{Rho}$ - VPS39 siRNA vs VPS41 siRNA, 1 µM, 30 min | | -654.5 | -825.1 to -483.9 | 0.0124 | 44.71 | 1.038 |
| AV5.3$^{Rho}$ - RISC-free siRNA vs VPS8 siRNA, 1 µM, 30 min | | -2434 | -2832 to -2036 | 0.0002 | 18.16 | 3.422 |
| AV5.3$^{Rho}$ - RISC-free siRNA vs TGF-BRAP1 siRNA, 1 µM, 30 min | | -1584 | -1994 to -1174 | 0.0011 | 12.24 | 3.029 |
| AV5.3$^{Rho}$ - RISC-free siRNA vs VPS39 siRNA, 1 µM, 30 min | | -1900 | -2311 to -1489 | 0.0007 | 14.71 | 3.001 |
| AV5.3$^{Rho}$ - RISC-free siRNA vs VPS41 siRNA, 1 µM, 30 min | | -2555 | -2962 to -2147 | 0.0002 | 19.66 | 3.075 |
| AV5.3$^{Rho}$ - RISC-free siRNA vs VPS8 siRNA, 1 µM, 30 min | 3F | -220 | -282.1 to -158.0 | <0.0001 | 7.096 | 58.30 |
| AV5.3$^{Rho}$ - RISC-free siRNA vs TGF-BRAP1 siRNA, 1 µM, 30 min | | -275 | -334.6 to -215.3 | <0.0001 | 9.258 | 49.89 |
| AV5.3$^{Rho}$ - RISC-free siRNA vs VPS39 siRNA, 1 µM, 30 min | | -256.8 | -316.7 to -197.0 | <0.0001 | 8.614 | 50.66 |
| AV5.3$^{Rho}$ - RISC-free siRNA vs VPS39 siRNA, 1 µM, 30 min | | -260.3 | -321.0 to -199.5 | <0.0001 | 8.593 | 53.51 |

| | | | | | | |
|---|---|---|---|---|---|---|
| ZF5.3-DHFR$^{Rho}$ - 0.1 μM vs 0.5 μM, 30 min | 4B | 8847 | 8087 to 9607 | 0.0009 | 60.90 | 1.669 |
| ZF5.3-DHFR$^{Rho}$ - 0.5 μM vs 1.0 μM, 30 min | | 12959 | -10193 to 36111 | 0.0891 | 7.054 | 1.003 |
| AV5.3-DHFR$^{Rho}$ - 0.1 μM vs 0.5 μM, 30 min | | 11646 | 8743 to 14549 | 0.0007 | 11.99 | 3.382 |
| AV5.3-DHFR$^{Rho}$ - 0.5 μM vs 1.0 μM, 30 min | | 14758 | 75.82 to 29440 | 0.0495 | 3.963 | 2.204 |
| ZF5.3-DHFR$^{Rho}$ vs AV5.3-DHFR$^{Rho}$ - 0.1 μM, 1 hr | 4C | 35.66 | 12.13 to 59.18 | 0.0039 | 3.061 | 40.84 |
| ZF5.3-DHFR$^{Rho}$ vs AV5.3-DHFR$^{Rho}$ - 0.5 μM, 1 hr | | 2.874 | -56.24 to 61.98 | 0.9224 | 0.09791 | 45.61 |
| ZF5.3-DHFR$^{Rho}$ vs AV5.3-DHFR$^{Rho}$ - 1.0 μM, 1 hr | | -37.89 | -122.2 to 46.41 | 0.3736 | 0.8949 | 77.39 |
| AV5.3-DHFR$^{Rho}$ - RISC-free siRNA vs Lipofectamine control, 1 μM, 1 hr | 4D | -423.5 | -26161 to 25314 | 0.9369 | 0.09212 | 1.582 |
| AV5.3-DHFR$^{Rho}$ - RISC-free siRNA vs VPS39 siRNA, 1 μM, 1 hr | | -2757 | -16797 to 11284 | 0.4271 | 1.038 | 1.655 |
| AV5.3-DHFR$^{Rho}$ - RISC-free siRNA vs TGF-BRAP1 siRNA, 1 μM, 30 min | | 1201 | -11886 to 14288 | 0.7232 | 0.4095 | 1.927 |
| AV5.3-DHFR$^{Rho}$ - RISC-free siRNA vs VPS39 siRNA, 1 μM, 1 hr | 4E | -88.94 | -135.1 to -42.76 | 0.0003 | 3.841 | 69.68 |
| AV5.3-DHFR$^{Rho}$ - RISC-free siRNA vs TGF-BRAP1 siRNA, 1 μM, 30 min | | -92.57 | -133.9 to -51.29 | <0.0001 | 4.479 | 64.28 |
| hDHFR vs AV5.3-DHFR specific | 5C | -3.460 | -7.489 to 0.5693 | 0.0651 | 3.972 | 1.861 |

| | | | | | | |
|---|---|---|---|---|---|---|
| activity values | | | | | | |
| hDHFR vs ZF5.3-DHFR specific activity values | | -3.610 | -11.24 to 4.025 | 0.1106 | 5.099 | 1.075 |
| AV5.3-DHFR vs ZF5.3-DHFR specific activity values | | -0.1500 | -5.405 to 5.105 | 0.8239 | 0.2767 | 1.132 |
| AVCF vs ZFCF specific activity values | 4D | -1.520 | -1.851 to -1.189 | 0.0007 | 14.51 | 3.041 |
| AV5.3-DHFR$^{Rho}$ vs AV5.3-DHFR$^{Rho}$ + MTX - 0.1 μM, 1 hr | S7A | -882.4 | -2787 to 1023 | 0.2250 | 1.563 | 2.721 |
| AV5.3-DHFR$^{Rho}$ vs AV5.3-DHFR$^{Rho}$ + MTX - 0.5 μM, 1 hr | | -7191 | -10023 to -4359 | 0.0025 | 7.261 | 3.724 |
| ZF5.3-DHFR$^{Rho}$ vs AV5.3-DHFR$^{Rho}$ + MTX - 1.0 μM, 1 hr | | -10027 | -25220 to 5167 | 0.1071 | 2.732 | 2.084 |
| Lys9$^{Rho}$ vs Lys9$^{Rho}$ + ZF5.3 | S4 | 1.643 | -0.1725 to 3.458 | 0.0749 | 1.825 | 42.99 |
| Lys9$^{Rho}$ + ZF5.3 vs Lys9$^{Rho}$ + AV5.3 | | 0.6727 | -1.344 to 2.679 | 0.5038 | 0.6733 | 50.29 |
| Lys9$^{Rho}$ vs Lys9$^{Rho}$ + RNAiMAX | | 40.60 | 32.11 to 59.08 | <0.0001 | 6.843 | 38.41 |
| AV5.3-DHFR$^{Rho}$ vs AV5.3-DHFR$^{Rho}$ + MTX - 0.1 μM, 1 hr | S7B | -5.357 | -18.30 to 7.583 | 0.4137 | 0.8205 | 109.6 |
| AV5.3-DHFR$^{Rho}$ vs AV5.3-DHFR$^{Rho}$ + MTX - 0.5 μM, 1 hr | | -122.9 | -150.1 to -95.66 | <0.0001 | 8.911 | 163.7 |
| ZF5.3-DHFR$^{Rho}$ vs AV5.3-DHFR$^{Rho}$ + MTX - 1.0 μM, 1 hr | | -180.2 | -256.1 to -104.2 | <0.0001 | 4.749 | 59.09 |
| hDHFR pH 7.5 vs hDHFR pH 6.5 specific activity values | S8E | -0.4450 | -9.813 to 8.923 | 0.6627 | 0.5834 | 1.015 |
| hDHFR pH 7.5 vs | | -2.540 | -5.943 to 0.8627 | 0.0685 | 7.897 | 1.085 |

| | | | | | | |
|---|---|---|---|---|---|---|
| hDHFR pH 5.5 specific activity values | | | | | | |
| hDHFR pH 7.5 vs hDHFR pH 4.5 specific activity values | | -6.985 | -8.642 to -5.328 | 0.0086 | 34.78 | 1.231 |

#### XIII. Supplementary Figures 1-12

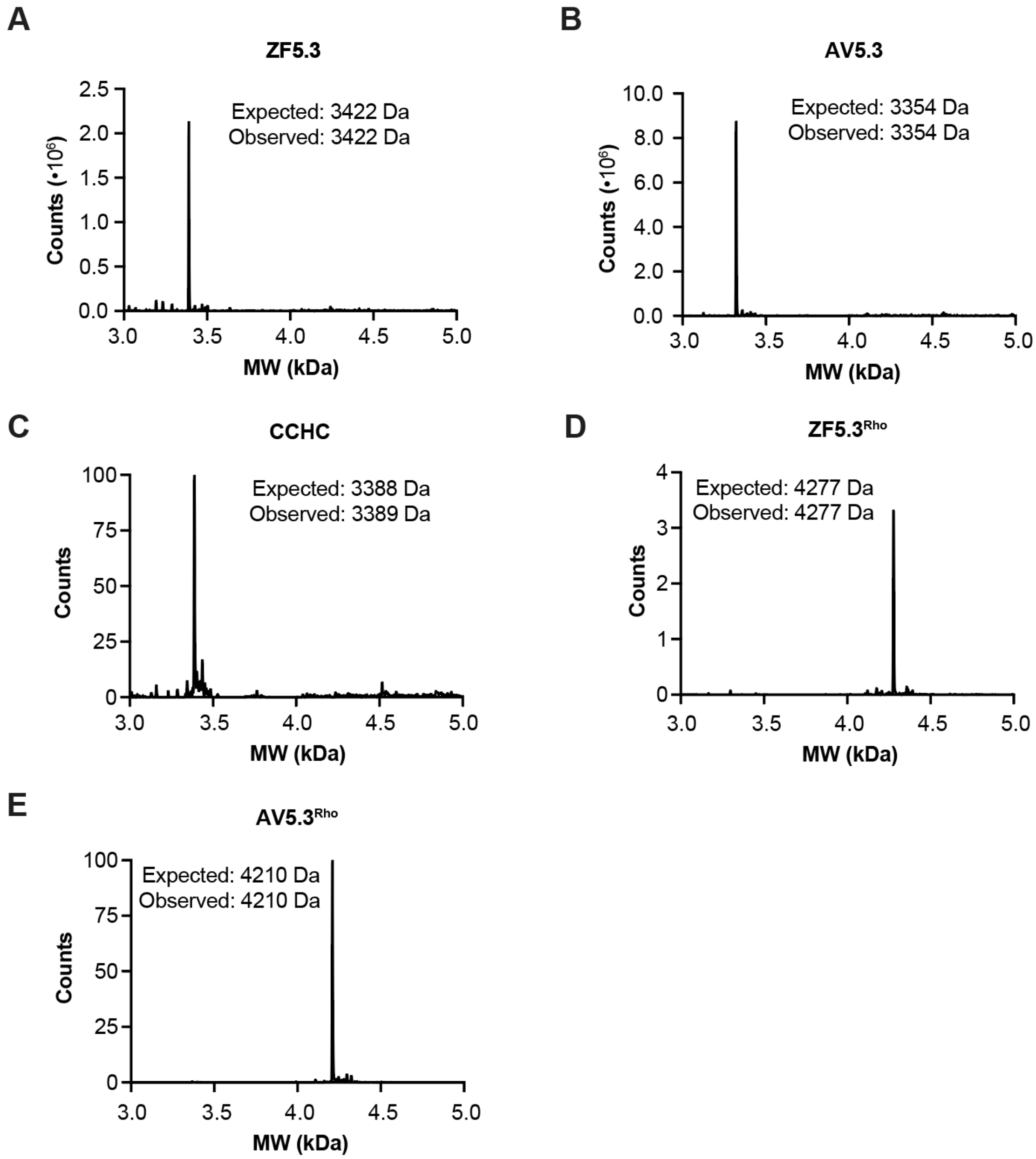


**Figure S1.** LC-MS analysis of purified ZF5.3, CCHC, and AV5.3 and Rho-labeled derivatives. Deconvoluted mass spectra of **(A)** ZF5.3; **(B)** AV5.3; **(C)** CCHC; **(D)** ZF5.3$^{Rho}$; **(E)** AV5.3$^{Rho}$.

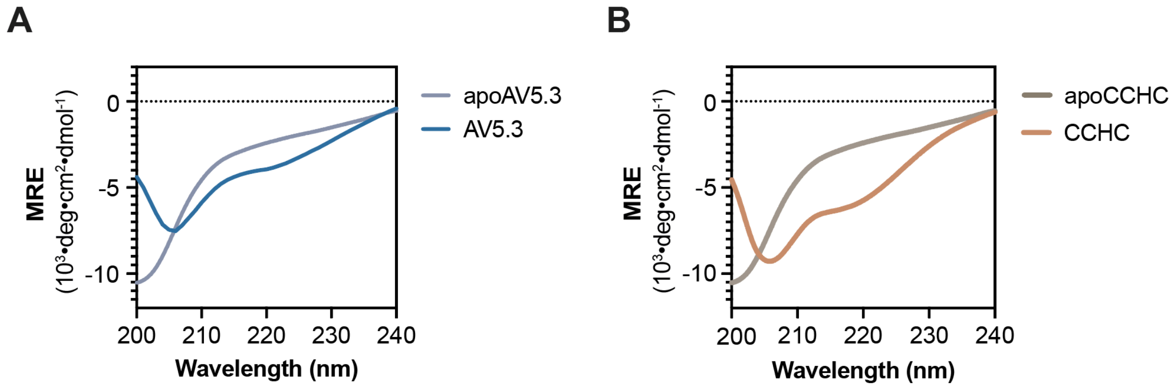


**Figure S2.** Wavelength-dependent CD spectra of apo and Zn(II)-coordinated **(A)** AV5.3 and **(B)** CCHC at 115 μM concentration in a buffer containing 20 mM Tris-HCl, 150 mM KCl, pH 7.5. Measurements were collected before and after the addition of 2 molar equivalents of Zn(II). For further details, refer to **Supplemental Methods**.

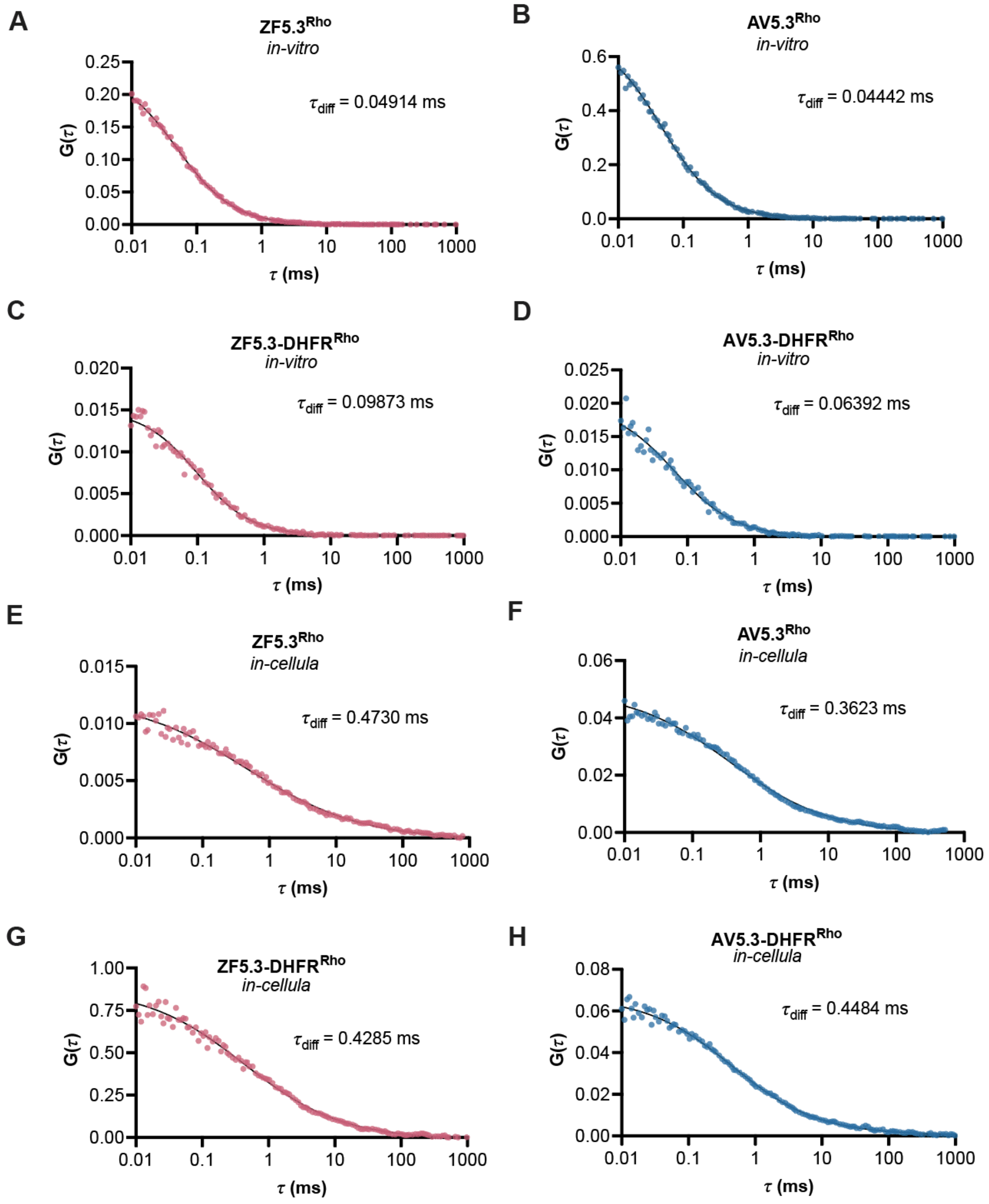


**Figure S3** Autocorrelation traces generated during FCS analysis of ZF5.3$^{Rho}$, AV5.3$^{Rho}$, ZF5.3-DHFR$^{Rho}$, and AV5.3-DHFR$^{Rho}$ in vitro and in Saos-2 cells. (A-D) Autocorrelation traces generated for *in-vitro* samples of 100-500 nM of ZF5.3$^{Rho}$, AV5.3$^{Rho}$, ZF5.3-DHFR$^{Rho}$, and AV5.3-DHFR$^{Rho}$ in DMEM cell media are shown. All traces were fitted to a 3D diffusion model (solid dark curve) to obtain *in-vitro* diffusion times ($\tau_{diff}$) as described in Methods. (E-H) Representative autocorrelation traces of Saos-2 cells treated with 500 nM ZF5.3$^{Rho}$, AV5.3$^{Rho}$, ZF5.3-DHFR$^{Rho}$, and AV5.3-DHFR$^{Rho}$ for 30-60 min. Traces were fitted using a 3D anomalous diffusion model as previously described3 to obtain *in-cellula* diffusion times. *In-cellula* diffusion times are longer than those *in-vitro* due to increased viscosity and impaired diffusion in the cytosol.

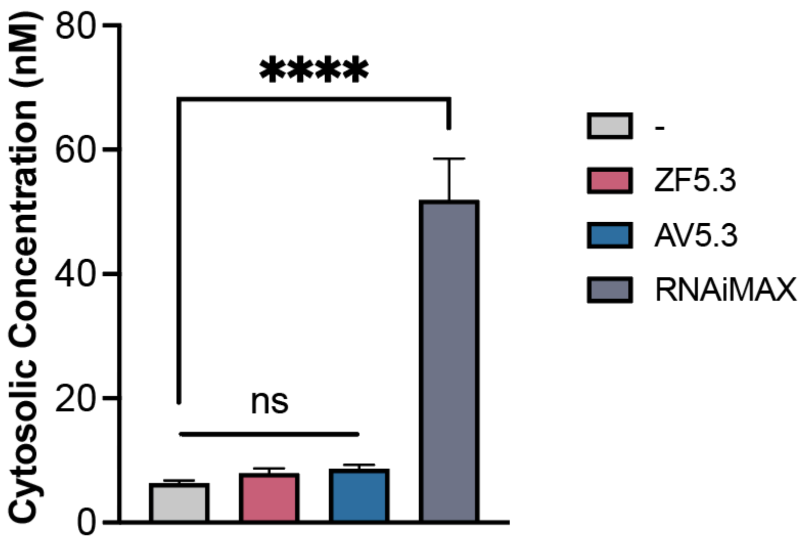


**Figure S4:** Codelivery of AV5.3 and Lys9$^{Rho}$ results in no detectable leakage of Lys9$^{Rho}$ into cytosol. Saos-2 cells were treated with 1 μM Lys9$^{Rho}$ plus either lipofectamine RNAiMAX (16 μL/mL), ZF5.3 (1 μM) or AV5.3 (1 μM), incubated for XX min, and the endosomal escape of Lys9$^{Rho}$ determined using FCS as described in **Methods**. Error bars represent the standard error of the mean. ****$p < 0.0001$, and not significant (ns) for $p > 0.05$ from one-way ANOVA with Dunnett post-test.

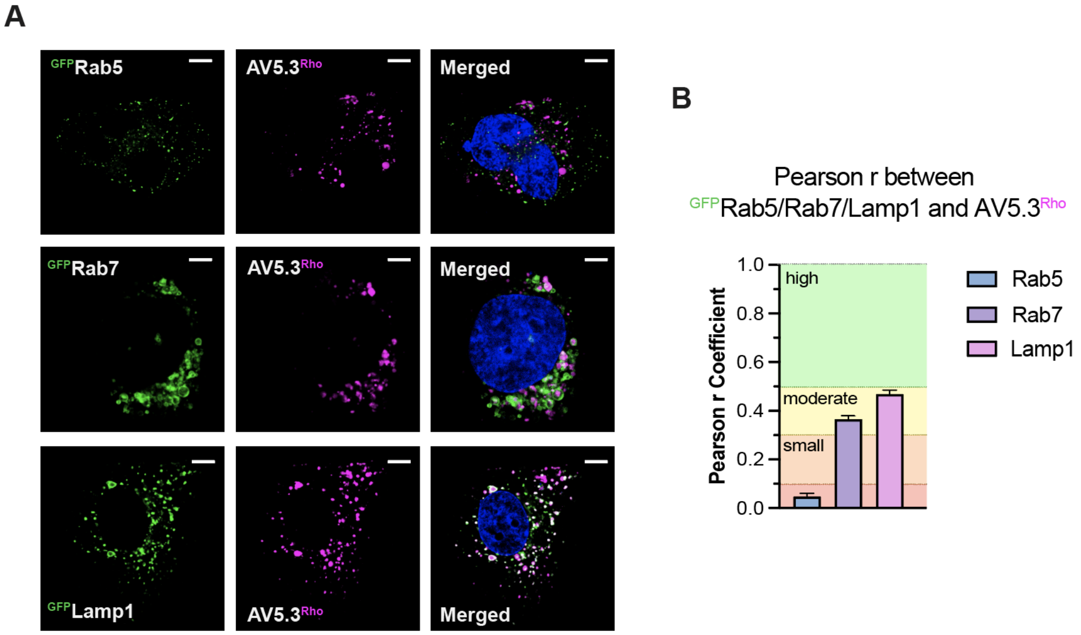


**Figure S5:** AV5.3$^{Rho}$ accumulates in Rab7+ and Lamp1+ endosomes. (A) Representative live-cell confocal microscopy images of Saos-2 cells expressing Rab5-, Rab7-, or Lamp1-GFP and incubated with AV5.3$^{Rho}$ (1 μM) for 30 min then with Hoechst 33342 as a counterstain (300 nM) for 5 min. Cells were lifted using TrypLE Express to remove exogenously bound materials and replated for microscopy as described in Methods. Scale bars: 5 μm. (B) Pearson correlation coefficients representing the co-localization of each GFP marker with AV5.3$^{Rho}$. Error bars represent the standard error of the mean.

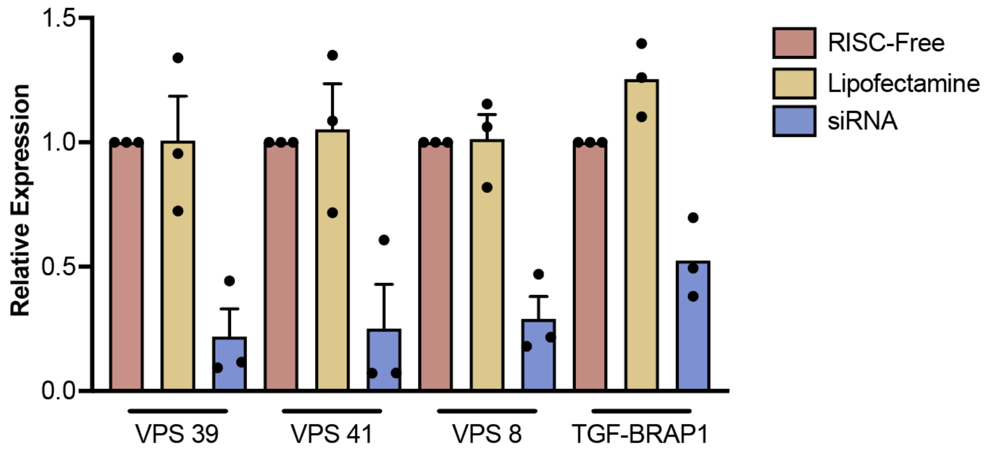


**Figure S6.** qPCR analysis of siRNA knockdowns. The gene expression levels of HOPS complex subunits (VPS39 and VPS41) and CORVET subunits (VPS8 and TGF-BRAP1) were determined using RT-qPCR and compared to RISC-free and lipofectamine only negative controls. Data are represented as mean ± SEM with three biological replicates.

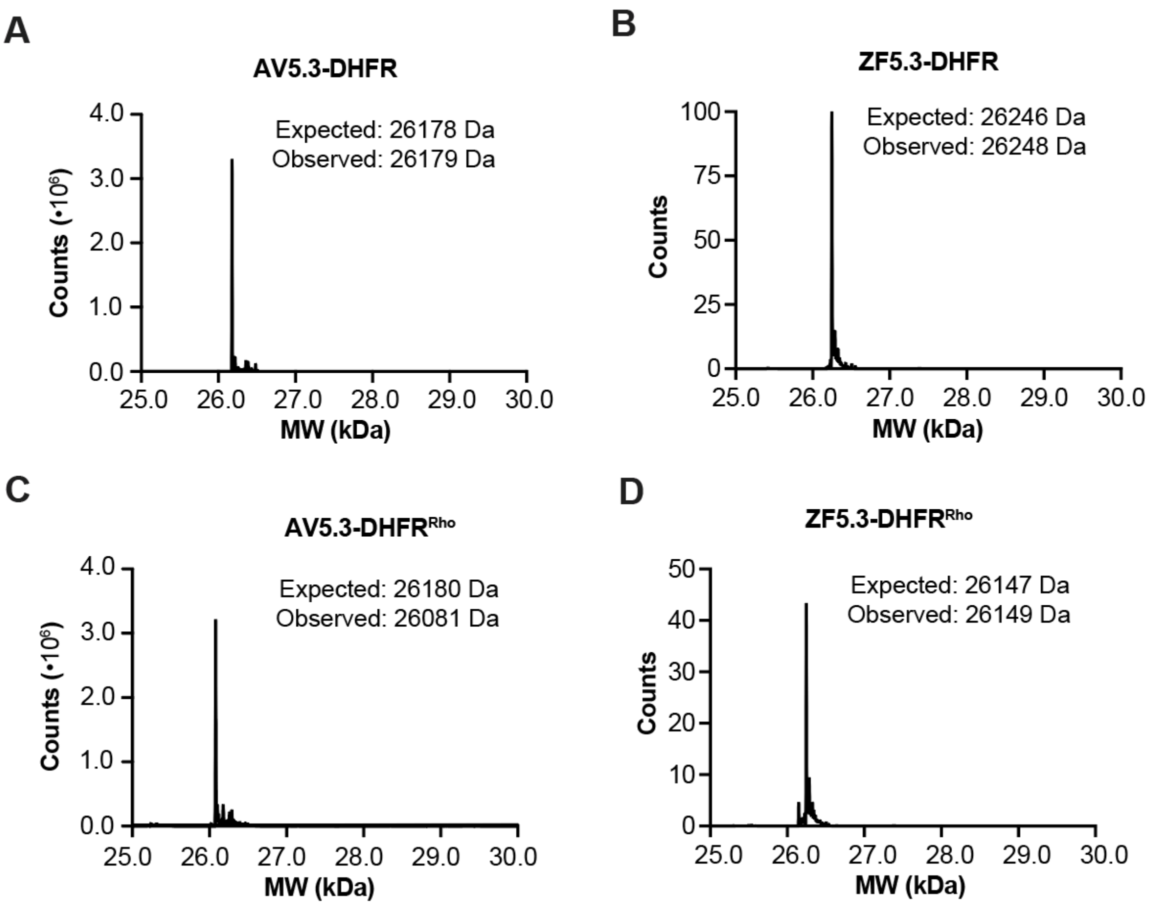


**Figure S7.** LC-MS analysis showing the deconvoluted mass spectra of purified AV5.3-DHFR, ZF5.3-DHFR and Rho-labeled derivatives. Deconvoluted spectra of **(A)** AV5.3-DHFR; **(B)** ZF5.3-DHFR; **(C)** AV5.3-DHFR$^{Rho}$; and **(D)** ZF5.3-DHFR$^{Rho}$.

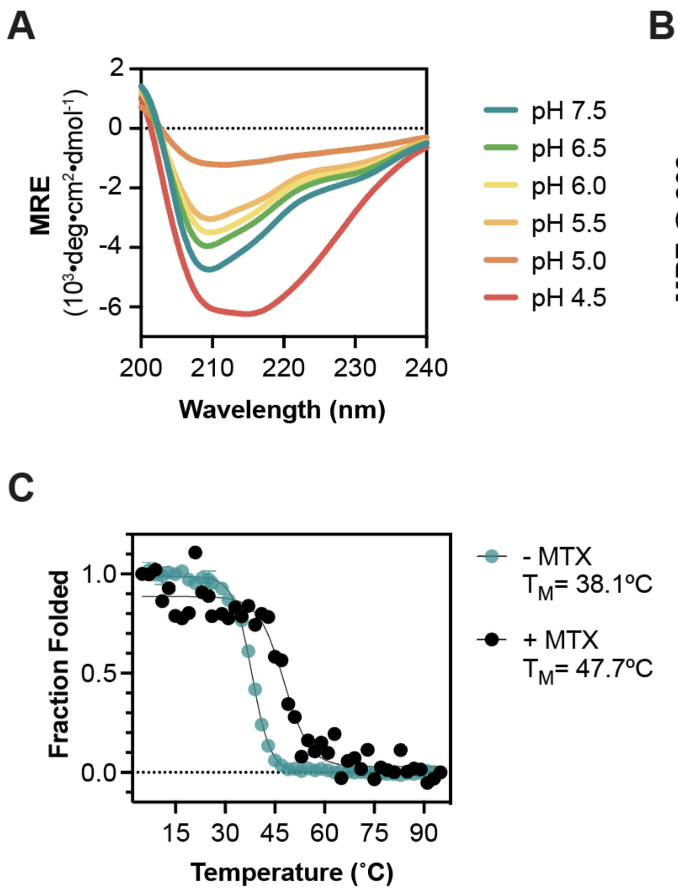


**Figure S8.** Wavelength-dependent CD spectra and temperature melts of 20 µM of AV5.3-DHFR at varying pH values or stabilized by MTX. (A) Wavelength CD spectra of AV5.3-DHFR showing a peak at 208 nm at pH 7.5 to 5.5. (B) Temperature melt of AV5.3-DHFR identifying a $T_M$ of 38.1˚C at pH 7.5, $T_M$ of 35.9˚C at pH 6.5, $T_M$ of 34.5˚C at pH 6.0, $T_M$ of 32.5˚C at pH 5.5. (C) Fraction folded of AV5.3-DHFR at pH 7.5 with or without being preincubated with 1 equivalence of MTX. The preincubation with MTX stabilized AV5.3-DHFR by 9.6˚C.

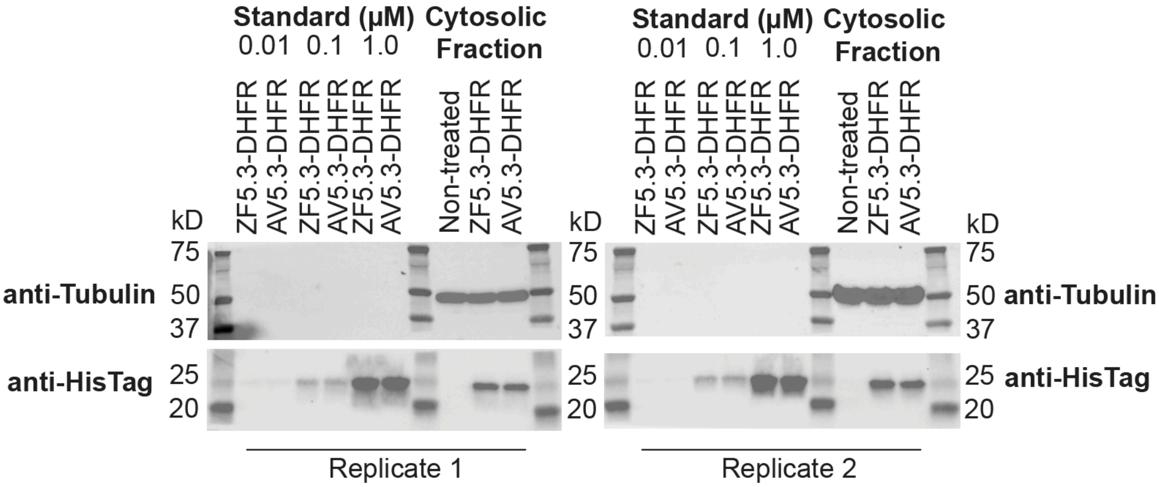


**Figure S9.** AV5.3-DHFR is fully intact when isolated from the cytosol of treated Saos-2 cells. Western blot analysis of cytosolic fractions isolated from Saos-2 cells treated with clear McCoy's media alone (non-treated) or 1 µM ZF5.3-DHFR or AV5.3-DHFR for 1 h. After treatment, cells were lysed, the cytosol was isolated by ultracentrifugation at 350 kg for 1 h, and the cytosolic fractions resolved using SDS-PAGE. After transfer to PVDF membranes, the blots were probed using an anti-tubulin antibody to identify cytosolic fractions and an anti-HisTag antibody to identify AV5.3-DHFR and ZF5.3-DHFR. Standards shown represent authentic samples of ZF5.3-DHFR and AV5.3-DHFR at the indicated concentration. Both the authentic samples and the materials isolated from the cytosol co-migrate with the 25 kD marker with no evidence of truncated fragments. Gel results are representative of two biological replicates.

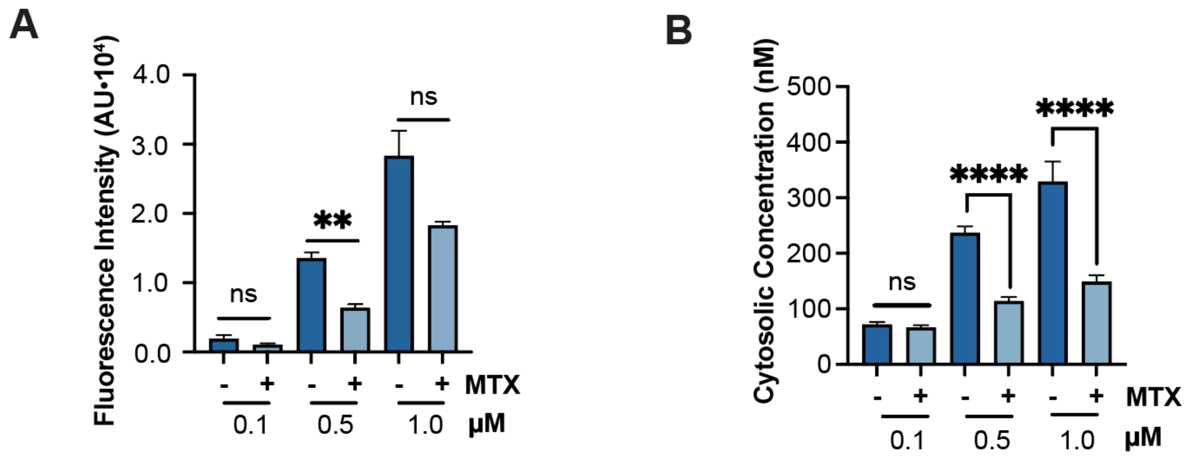


**Figure S10.** Effect of MTX on the total uptake and cytosolic delivery of AV5.3-DHFR$^{Rho}$. (A)Plot illustrating the effect of MTX on overall uptake and (B)cytosolic delivery of AV5.3-DHFR$^{Rho}$. The protein was preincubated with 1 equivalent of MTX for 30 min and then added to cells at a total treatment concentration of 0.1 − 1 μM for 1h. Cells were then trypsinized and analyzed by FC or FCS as described above. FC values are provided as median fluorescence intensity for the lissamine rhodamine B channel (MFI); n =20000 per condition in total with at least two biological replicates each (mean ± SEM). FCS values are provided in nM; n >20 for each FCS condition comprising at least three biological replicates each (mean ± SEM). Statistical significance comparing the given concentrations was assessed using the Brown−Forsythe and Welch one-way analysis of variance (ANOVA) followed by an unpaired t test with Welch's correction. ****$p \leq 0.0001$, ***$p \leq 0.001$, **$p \leq 0.01$, *$p \leq 0.05$.

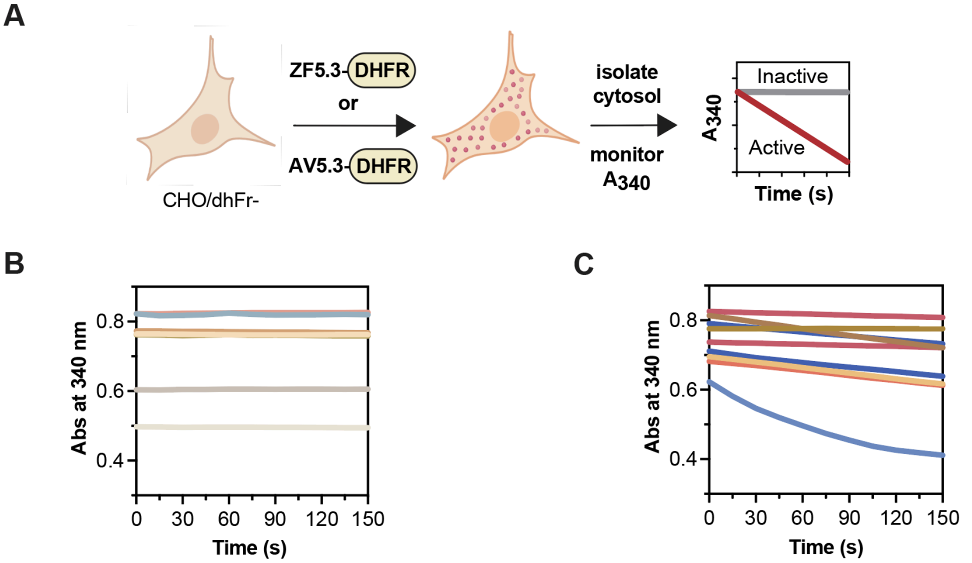


| Sample | hDHFR | hDHFR | hDHFR | hDHFR | AV5.3-DHFR | AV5.3-DHFR | ZF5.3-DHFR | ZF5.3-DHFR | NTCF | NTCF | NTCF + hDHFR | NTCF + hDHFR | AVCF | ZFCF |
|---|---|---|---|---|---|---|---|---|---|---|---|---|---|---|
| Color ID | | | | | | | | | | | | | | |
| DHF | - | + | + | + | + | + | + | + | + | + | + | + | + | + |
| NADPH | + | - | + | + | + | + | + | + | + | + | + | + | + | + |
| MTX | - | - | - | + | - | + | - | + | - | + | - | + | - | - |

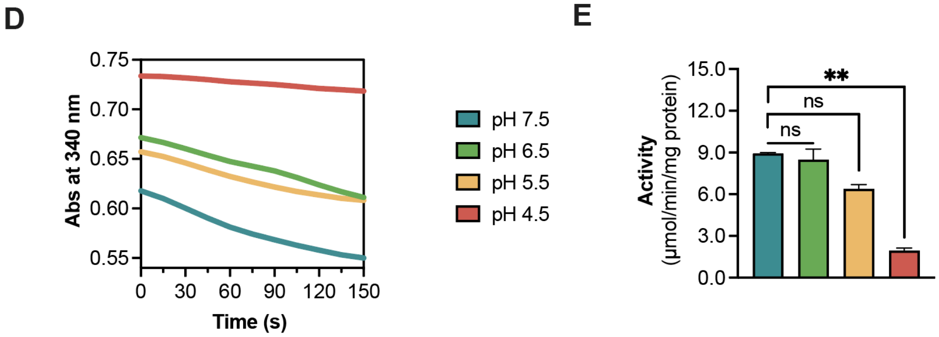


**Figure S11**. DHFR proteins activity assay curves. DHFR catalyzes the reduction of 7,8-dihydrofolate (DHF) to 5,6,7,8-tetrahydrofolate (THF) using a single equivalent of NADPH as a cofactor. The in vitro activities of AV5.3-DHFR, ZF5.3-DHFR, and human DHFR were measured spectrophotometrically by monitoring the decrease in NADPH absorbance at 340 nm in a reaction mixture supplemented with 250 nM enzyme and 50 μM DHF. (A) Post-delivery activity assay scheme. DHFR proteins were delivered to a DHFR-deficient cell line, CHO/dhFR-. The treated-cells cytosol was fractionated and subsequently quantified for DHFR activity by measuring the loss of $A_{340}$ signal over time. By taking into account the protein concentration, volume of protein added and the extinction coefficient of DHFR, we obtain a value of specific activity in mole/min/mg protein. (B) Control experiments where no change in Absorbance at 340 nm is observed. (C) Experimental results where loss of absorbance at 340 nm represents DHFR activity. (D) The activity of human DHFR is sensitive to pH. Specific activity values of hDHFR after being exposed to varying pHs. (E) Experimental results of DHF conversion to DHF by hDHFR samples at varying pHs.

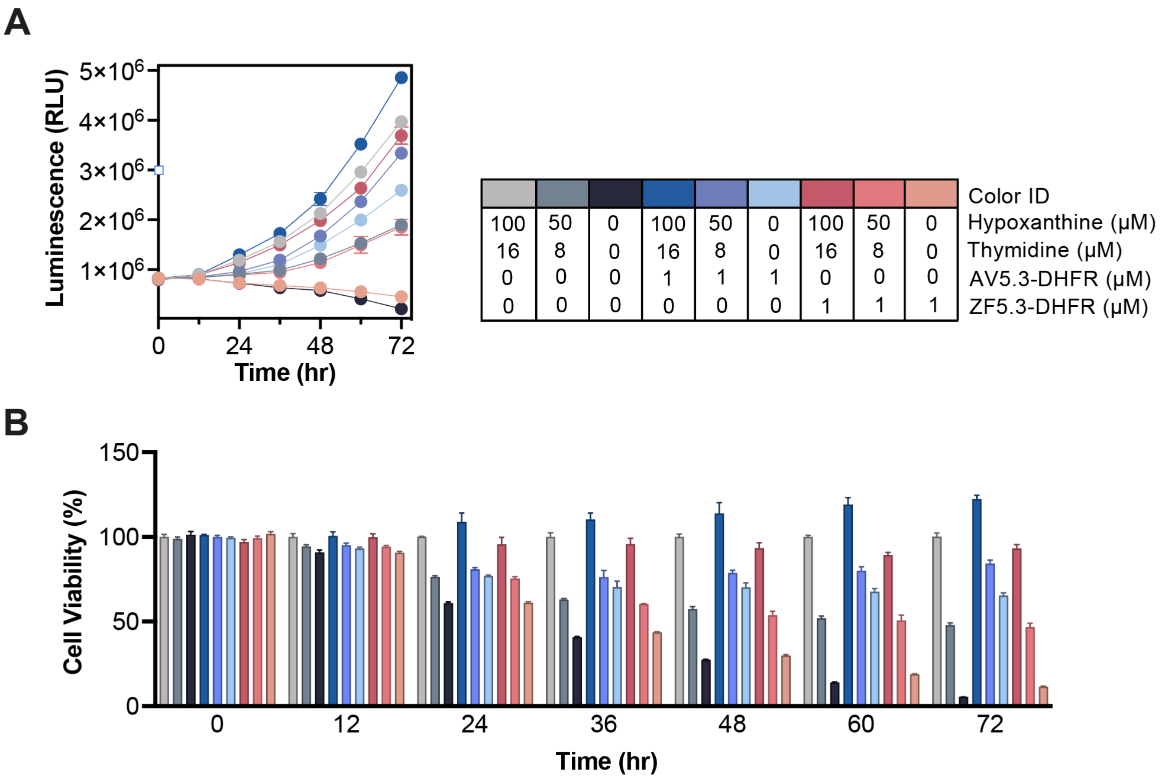


**Figure S12.** DHFR protein rescue in CHO/dhFr- cells. (A) Raw luminescence curves of ATP-activity in 5x10$^3$ CHO/dhFr- cells treated with varying concentrations of hypoxanthine/thymidine (HT) supplement and AV5.3-DHFR or ZF5.3-DHFR. (B) Percentage of viable cells quantified every 12 hrs for 72 hrs.
